# West Nile Virus spread in Europe - phylogeographic pattern analysis and key drivers

**DOI:** 10.1101/2022.11.10.515886

**Authors:** Lu Lu, Feifei Zhang, Bas B. Oude Munnink, Emmanuelle Munger, Reina S. Sikkema, Styliani Pappa, Katerina Tsioka, Alessandro Sinigaglia, Emanuela Dal Molin, Barbara B. Shih, Anne Günther, Anne Pohlmann, Martin Beer, Rachel A. Taylor, Frederic Bartumeus, Mark Woolhouse, Frank M. Aarestrup, Luisa Barzon, Anna Papa, Samantha Lycett, Marion P. G. Koopmans

## Abstract

Spread and emergence of West Nile virus (WNV) in Europe have been very different from those observed in North America. Here, we describe key drivers by combining viral genome sequences with epidemiological data and possible factors of spread into phylodynamic models. WNV in Europe has greater lineage diversity than other regions of the world, suggesting repeated introductions and local amplification. Among the six lineages found in Europe, WNV-2a is predominant, has spread to at least 14 countries and evolved into two major co-circulating clusters (A and B). Both of these seem to originate from regions of Central Europe. Viruses of Cluster A emerged earlier and have spread towards the west of Europe with higher genetic diversity. Amongst multiple drivers, high agriculture activities were associated with both spread direction and velocity. Our study suggests future surveillance activities should be strengthened in Central Europe and Southeast European countries, and enhanced monitoring should be targeted to areas with high agriculture activities.

## Introduction

Mosquito-borne viruses represent a considerable public health problem worldwide, causing infections in both humans and animals^1^. For the European region, West Nile virus (WNV) is one of the mosquito-borne viruses which can cause severe disease and death in humans and has been increasing in prevalence and geographic range over the past decade ^2^. WNV belongs to the family *Flaviviridae* (genus *Flavivirus*) with an enveloped, single-stranded RNA genome ^3^. The transmission cycle of WNV involves mosquitos (mainly of the *Culex* species) as vectors and birds as amplifying reservoir hosts^4^, while humans and other mammals are considered dead-end hosts^1^. Dead-end hosts are not thought to contribute significantly to transmission in the natural life cycle of the virus. However, for humans, the potential for virus transmission through blood transfusion and organ transplantation has impacted blood and transplantation donor programs, with mandatory screening for regions where exposure to WNV is possible. Currently nine distinct lineages (WNV-1 to WNV-9) of WNV have been identified globally, but little is known about their phenotypic properties^5^. WNV-1 and WNV-2 strains have been identified most frequently in human and animal cases in multiple continents, while strains within WNV-3 to WNV-9 have been detected from mosquitos, birds and equines sporadically in parts of Europe, Asia, and Africa^6^. The dominant lineage in Europe in recent years is WNV-2, although cases of WNV-1 have been reported^7,8^.

WNV circulation in Europe was first reported in the 1960s^9^. Since 1996, an increasing number of WNV outbreaks in humans and equines were detected in Southeast and Central Europe^10^. In past decades, WNV outbreaks in humans and animals have been found almost annually in previously non-endemic areas^11^. As surveillance and clinical testing of WNV infections are patchy, it is difficult to compare case notifications across Europe. Nevertheless, considerable differences have been observed between successive years, with for instance particularly severe regional WNV outbreak involving both humans and equids in 2018^12^ detected in Italy, Serbia, and Greece^13^. Since the beginning of the 2022 transmission season, over 60 outbreaks among birds (mainly in Italy) have been reported and over 800 human cases of WNV infections were reported in seven countries, mainly in Greece and Italy^7^. Unlike in the United States, where the initial WNV (WNV-1) incursion in 1999 led to rapid dispersal of the virus throughout the continent ^14^, patterns of spread have remained somewhat patchy in Europe and involved both WNV-1 and WNV-2. Although there have been studies on the presence and the spread of WNV in different European countries^15–17^, the overall dispersal history and the important drivers have yet to be determined. Therefore, we explored the possible added value of integrated analysis of different types of publicly available and newly acquired data to understand and potentially predict the trajectory of WNV dispersal across Europe.

To infer the virus dispersal history from sequence and other data, it is important to model how viruses are dispersed through space, between species or host-types. For inference of transmission patterns of viruses between discrete locations, a phylogenetic discrete traits analysis can be used^18^. If the dispersal history of viruses is in a continuous space setting^19^, the coordinates of the ancestral nodes of the tree could be inferred from the sampled locations and two-dimensional diffusion rate^19^. The inferred transmission pattern between locations and the spreading rate can be further modelled as functions of a combination of spatial environmental factors, and the contribution of the factors modulating the transmission pattern can be determined^20^. In addition, the importance of factors which may affect the viral effective population size (a measure of the extend of circulation inferred from sequence diversity) over time can be estimated using a Generalized Linear Model (GLM) approach based on the temporal data (for example, seasonal or climate change signals)^21^. In addition, increased virus transmission rates and dispersal are signals of changing outbreak potential, although drivers for spread during outbreaks and drivers of incursion to new regions might differ.

In this study, we explored the dispersal history of WNV in Europe and the underlying drivers in regions with yearly WNV outbreaks and in regions with sporadic outbreaks. We used phylodynamic models which incorporated: (i) sequencing data (Supporting file 1), (ii) epidemiological data (host species, sampling time and coordinates, as well as travel history of humans if available) (Supporting file 1), (iii) socio-economic data (population and GDP), and (iv) environmental data (climatic, land cover, land use, biodiversity) (Supporting file 2 and 3). Updated sequencing data and epidemiology data were provided by a European collaborative consortium initiated by a group of experts across Europe (https://www.veo-europe.eu/). Specifically, we described the evolution and genetic diversity of WNV in Europe; we then explored the pattern of transmission between and within countries and the possible predictors of spread.

## Results

### Diverged WNV lineages found in Europe

We found a positive correlation between cumulative human cases in EU/EEA reported by ECDC (between 2008-2021, total n=4188) and the number of WNV-2 sequences (between 2004-2021, total n=485) provided from different countries (R= 0.75, p<0.005), despite the varied sequencing capacity among countries (Figure S1a). Among the 22 European countries that have human cases reported, 15 of them have sequences available, including Greece, Italy, Romania, Serbia, Hungary, Spain, Austria, France, Bulgaria, Germany, Netherlands, Slovenia, Portugal, Slovakia and Czechia; sequences are missing from six countries (Croatia, Kosovo, Albania, Bosnia and Herzegovina, North Macedonia, Montenegro, and Cyprus) in the east region (Figure S1b). By comparing the number of sequences available versus the number of human cases reported, the sequencing effort in central and north regions are better than the south and east regions (Figure S1b).

The maximum likelihood tree of all available WNV genome sequences showed that six lineages (out of the total 9 lineages globally^22^) were detected in Europe^6^ (Figure 1). Among the six lineages, WNV-2 had the largest number of sequences available, accounting for 82% of all WNV sequences detected in Europe so far, and the widest diffusion, since it has been found in 15 European countries (Figure S2). The earliest WNV-2 genome (JX041631.1) was sampled from birds in Eastern Europe (Ukraine) in 1980 (Figure 1b and 1c). The dominant sub-lineage 2a (WNV-2a) emerged in 2004 and became the dominant lineage in the past ten years, while the other lineages were rarely seen in the same time period. In addition, there was a separate small sub-lineage 2b (WNV-2b) composed of sequences mainly from Romania, Italy and Russia in 2011-2015, and one detection in Greece in 2018^23^ (Figure 1b and S2).

**Figure 1.**
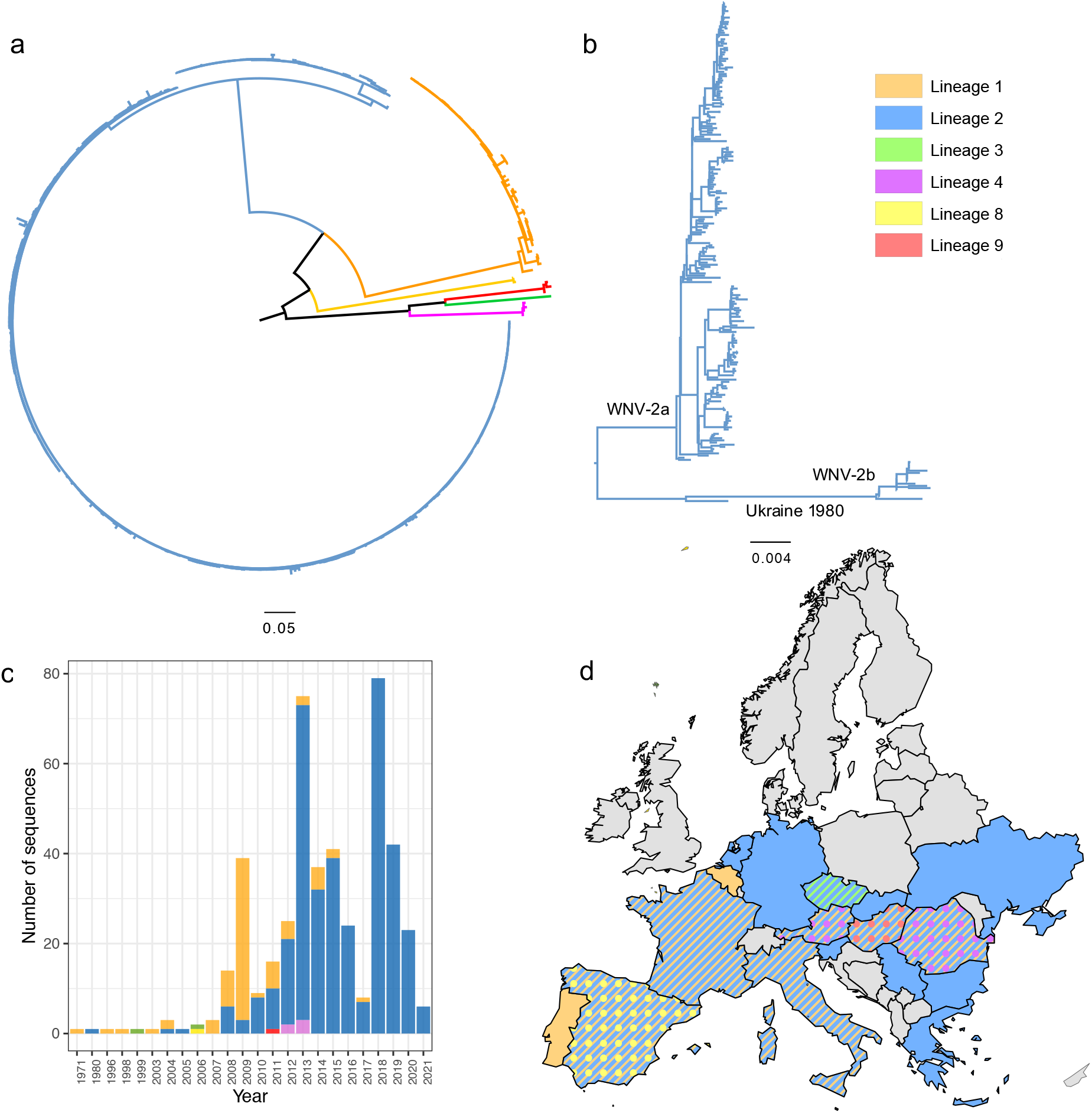
Phylogenetic analysis of WNV full and partial nucleotide sequences detected from Europe. The evolutionary distances were computed using the optimal GTR+I model, the phylogenetic tree was constructed with the Maximum likelihood method. Bootstrap values are given for 1000 replicates. (a) ML tree of all lineages found in Europe; the branches of the tree are colored by lineages; (b) The subtree of WNV-2 sequences; (c) The WNV lineages distribution over time using the same color showing on the tree; (d) The geographical distribution of WNV lineages, using the same lineage color showing on the tree, and differentiate by shape if multiple lineages were found within the same country.

In comparison, sequences of the second largest lineage (WNV-1) have been found in seven European countries (Austria, Italy, Spain, France, Hungary, Romania and Portugal) since 1971. Most WNV-1 sequences were reported in Italy (72% of total WNV-1 sequences), including all currently available WNV-1 sequences from humans. WNV sequences belonging to lineages 3, 4, 8 and 9 were only sporadically reported and all of them were collected from mosquitos: WNV-3 strains were only found in Czech Republic in 1997 and 2006; WNV-4 in Romania in 2012-13; WNV-8 were only found in Spain in 2006, while WNV-9 genomes were obtained in Austria in 2013 and Hungary in 2011 (Figure 1c,1d and Figure S2). In addition, up to 2021, these lineages (WNV-1, 3, 4, 8 and 9) were only collected from non-human hosts (mainly birds or mosquitos, very few equines), except 9 WNV-1 genomes from humans in Italy between 2009-2013.

### Phylodynamics of the predominant WNV-2a

Considering the distinct genetic diversity, the number of sequences available, and the dominance of WNV-2 as cause of human disease, the detailed phylodynamic analyses were focused on WNV-2a. These sequences were collected from 14 countries between 2004 to 2021, with most available sequences from Greece (n=64), Italy (n=50), and Germany (n=46) (Supporting file 1). The sequences were collected from 6 host types, with 30% from bird, 30% from mosquito and 40% from humans and mammals (Figure S3 a and b).

The time-scaled phylogenies of WNV-2a in Europe rooted with the first genomes found in Hungary, with the estimated time to the most recent common ancestor (TMRCA) dated back to July 2003, with a 95% highest posterior density (HPD) interval between November 2002 and July 2004. The WNV-2a sequences in Europe were split into two distinct sub-clusters (A and B) on the tree composed of full genome sequences (Figure 1a), which were previously named the central eastern clade and southeastern clade, respectively (Figure 2a). We could see that the two clusters evolved independently after the incursion from central Europe. Both Cluster A and Cluster B are present in Serbia, Slovakia, Hungary, Slovenia and Italy. Cluster A has also been transmitted to Austria, Czech Republic, Germany, the Netherlands, France and Spain; while Cluster B has also been found in Greece, Romania and Bulgaria.

**Figure 2.**
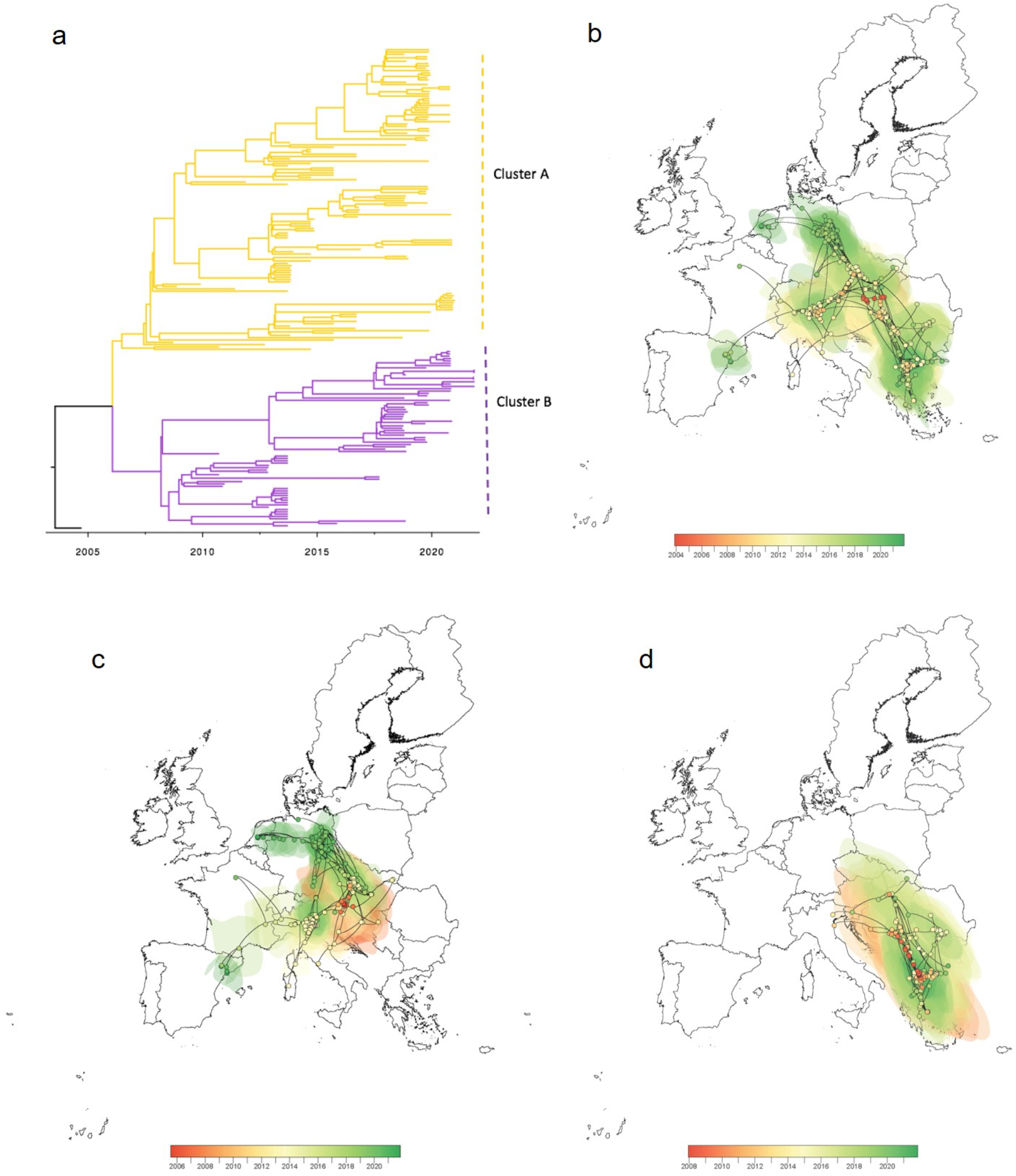
Time-scaled phylogeny of WNV-2a genomes in Europe. (a) Time-scaled MCC (maximum clade credibility) tree of WNV full genome sequences isolated in Europe (n=192), the two cluster A and B are labelled on the right. A distinct phylogeographic analysis has been based on Ns3 gene by continuous phylogeographic inference based on 1,000 posterior trees. Spatiotemporal diffusion of all WNV-2a in Europe (b), of cluster A (c) and Cluster B (d). These MCC trees are superimposed on 80% the highest posterior density (HPD) interval reflecting phylogeographic uncertainty. Nodes of the trees, as well as HPD regions, are colored by timescale from red (the time to the most recent common ancestor, TMRCA) to green (most recent sampling time), and oldest nodes (and corresponding HPD regions) are here plotted on top of youngest nodes.

Next, we estimated the time of emergence and rate of evolution, as well as the population dynamics and speed of diffusion for the two WNV-2a clusters A and B. WNV of Cluster A emerged in approximately July 2006 (with 95% HPD between January 2005 and March 2007; Figure 3a). The estimated ancestor country of Cluster A was Austria, with subsequent sequences identified mainly in Germany and Italy (Figure 2). With a similar evolution rate, Cluster B emerged later in June 2007 (with 95% HPD between March 2006 and December 2008) from Hungary, and most subsequent sequences were collected from Greece (Figure 2, 3a and 3b).

**Figure 3.**
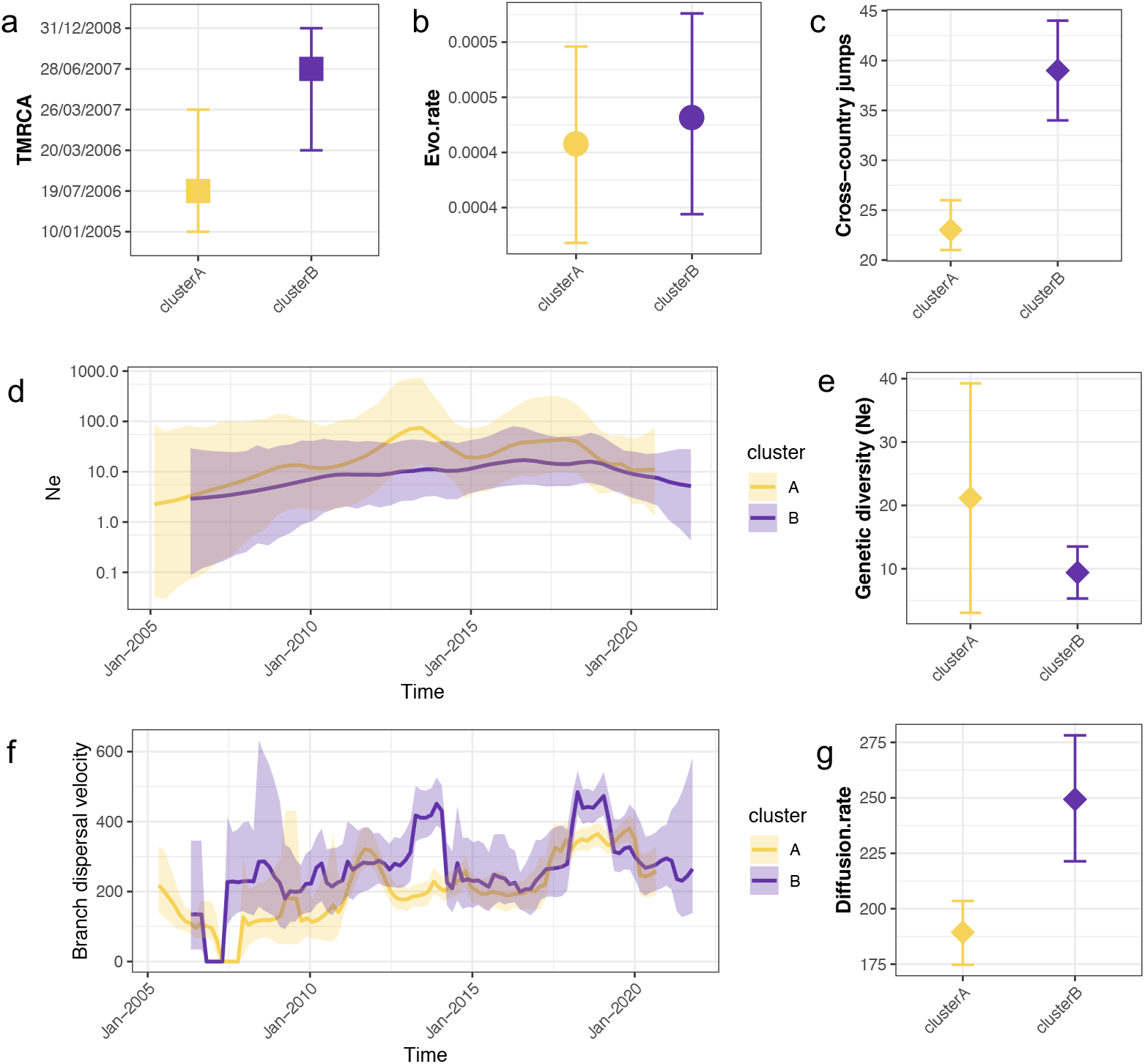
Comparisons of two co-circulating clusters of WNV-2a in Europe. (a) The mean time of the most recent ancestor (TMRCA) and 95%HPD interval for each cluster. (b) The mean clock rate and 95% HPD interval for each cluster. (c) Mean number of Markov jump between countries and 95%HPD interval for each cluster. (d) Estimation of effective population size via time and 95% HPD interval. The logarithmic effective number of infections (Ne) vs. viral generation time (t), representing effective transmissions is plotted over time. (e) Mean genetic diversity (Ne) and 95% HPD interval. (f) The weighted branch dispersal velocity (km/y) through time and 95% HPD interval. (g) The weighted mean diffusion rate (km/y) and 95% HPD interval.

We also compared the population dynamics of the two clusters. The effective population size (Ne) of cluster A peaked around 2014, which corresponded to the expansion phase of the epidemic, and then decreased slightly till the second peak at around 2018, which was consistent with a high WNV activity in multiple regions. In comparison, there were less important changes in Ne of Cluster B (Figure 3d). The higher genetic diversity translates into a higher mean Ne of cluster A compared to Cluster B. (Figure 3d, e).

WNV-2a spread at a high diffusion rate in Europe; the estimations vary between 88 km/y (full genome), 215 km/y (Ns3 gene) and 180 km/y (Ns5 gene), respectively. The dispersal velocity of Cluster B sequences was higher (mean of 249 km/y with 95% HPD between 221 and 278 km/y using Ns3 data) than those of Cluster A (mean of 189 km/y with 95% HPD between 175 and 203km/y using Ns3 data) (Figure 3f and 3g). This finding is consistent with the estimation of frequencies of between-countries transmissions: WNV of Cluster B tended to jump more frequently among countries, with an overall number of 39 (95% HPD between 34 and 44)— almost twice more often than Cluster A (Figure 3c).

### Quantified transmission frequencies between European countries and within Greece

We quantified frequencies of transmission among the 14 European countries where we have available WNV-2a sequences. We found transmissions were most likely to occur between neighboring countries. In addition, Austria and Hungary (both in Central Europe) had the most linkages with other countries (Figure 4 and Figure S5 and S6a).

**Figure 4.**
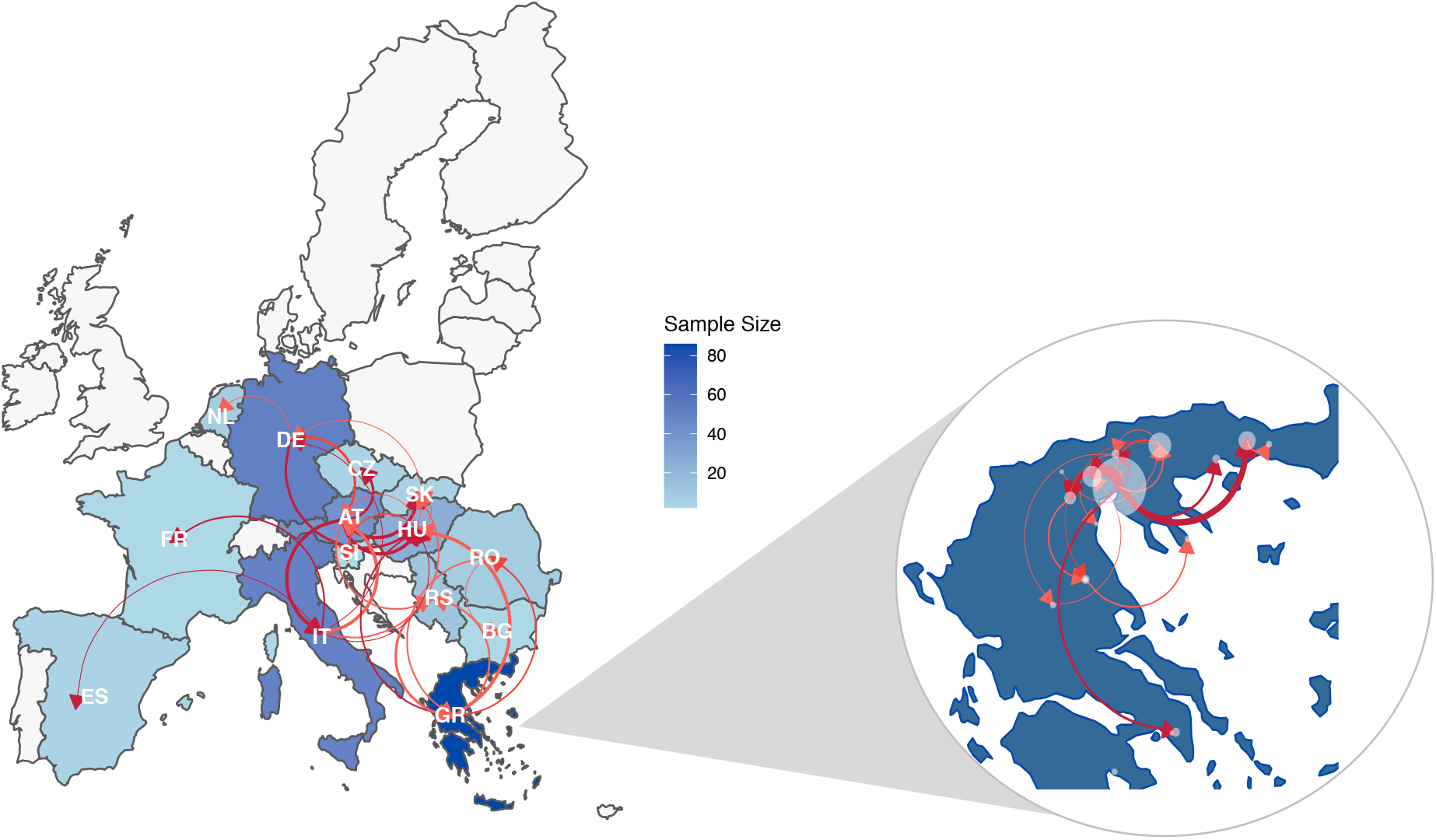
Quantified transmission network of WNV-2a between European countries and within Greece inferred using discrete trait models. Shape of colors on map indicates number of samples; edge weight indicates median number of transmissions between pairs of countries/regions; arrow on edge indicates transmission direction; color of edge from light to dark indicates Bayes Factor (BF) support from low to high only transmissions with BF >3 are shown).

Greece contributed the highest number of WNV-2a sequences in Europe. The time-scaled analysis showed Cluster B WNV has transmitted from central Europe (Hungary) to Greece and kept circulating within Greece almost annually except 2015-2016. In 2017, WNV-2a re-emerged in Greece and caused outbreaks in 2018 and continued to circulate during 2019–2021. Between 2017 and 2021, at least six separate introductions were observed (Figure S5). Three of them were from Greece only and fell in the same group with earlier isolates from Greece back to 2013. The other three groups included sequences from other countries, including Hungary, Romania and Bulgaria. Among them, sequences isolated from Hungary from 2018 to 2019 were most closely related to sequences from Greece sampled in the same time periods. This showed the possibility of both re-introduction from other countries and silent circulation maintained within Greece.

The discrete trait phylogeographic analysis estimated that approximately 19 transmission events between Greece and neighbouring countries occurred in the past decade. Countries that had the most frequent transmissions to Greece were Hungary (n=8), followed by Serbia (n=5) and Romania (n=4). The transmissions between Hungary and Greece occurred multiple times between 2010 and 2019, while the transmissions between Greece and other countries occurred mainly in 2012-2013. We further estimated the transmissions within Greece; between-region transmissions mainly occurred in north Greece and spread to the east and south regions (Figure 4b and Figure S6b).

### Continuous dispersal and impact of environmental factors on viral lineage dispersal

We reconstructed the dispersal history of WNV-2a in a continuous space (Movie S1, Figure 2 and 5) using a continuous phylogenetic model. WNV-2a was estimated to have originated from Central Europe in 2003-2004 (Hungary). One year later, WNV-2a dispersed to the eastern part of Austria and spread to the northwest and western regions (including Slovakia, Serbia, Hungary, Austria, Czech Republic, Slovenia, Germany, Netherlands, France and Spain) and southeastern countries (Greece, Romania, Italy, Bulgaria, Serbia, Slovakia, Slovenia). We could see that WNV emerged and circulated slowly in central Europe at first, and then spread to the southeast regions, and then in a westward direction. The frequency and velocity of diffusion was higher in 2013-14, then slowed down, and speeded up again around 2018 (Figure 3f). The periods with high diffusion rates were consistent with the timing when the re-emergence of WNV caused major outbreaks as well as the introduction to new geographic regions, e.g., further up north spreading of WNV-2a (Cluster A) towards Germany and the Netherlands.

**Figure 5.**
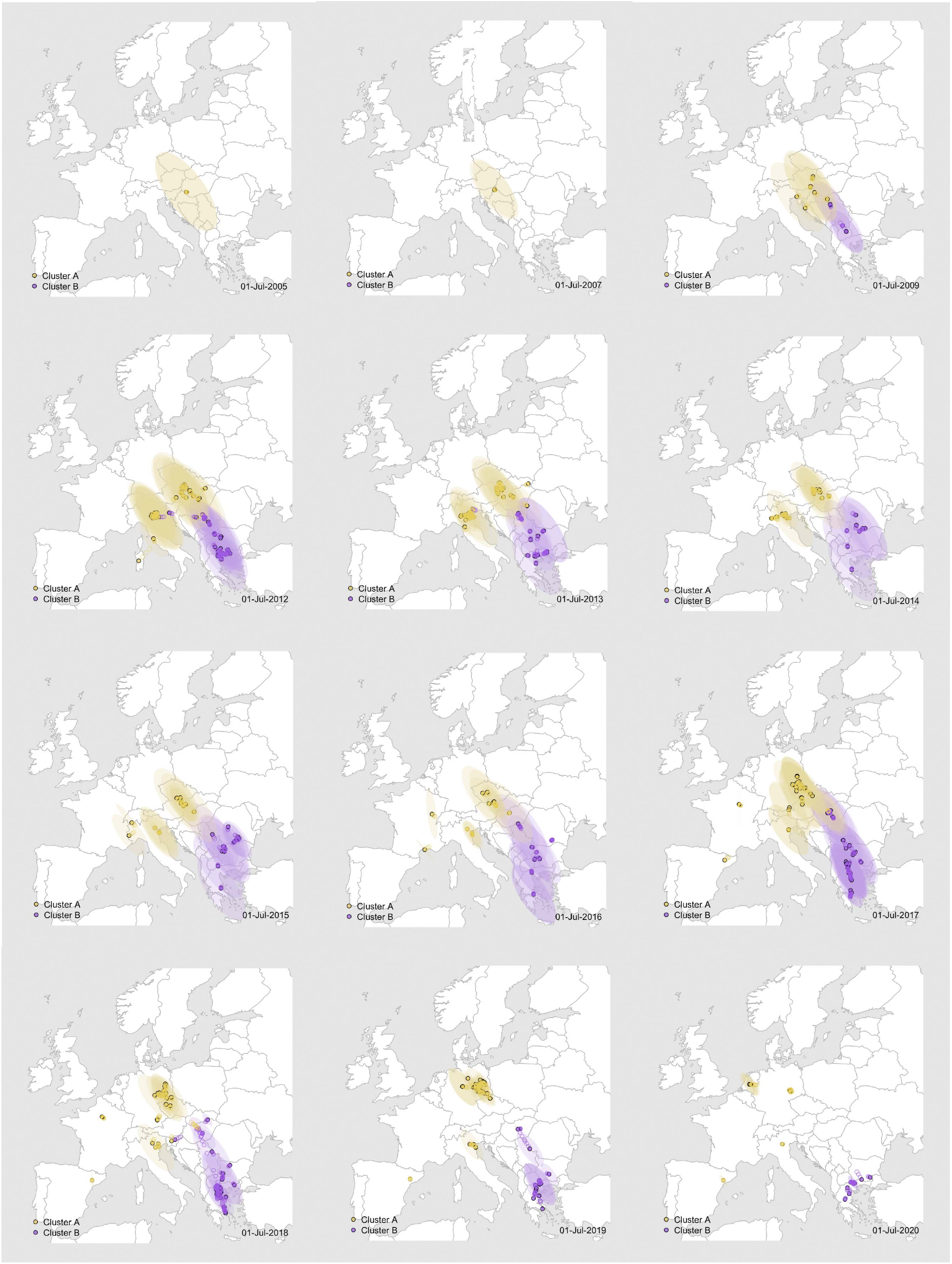
Dispersal history of WNV-2a in Europe between 2004 to 2021. Colors of the dots represent interpolated maximum clade credibility phylogeny positions for cluster A (yellow) and B (purple) from Ns3. Please see Movie S1 for the full movie.

We could also extract the spatio-temporal information embedded in the tree and test the correlation of the phylodynamic dispersion with gridded predictors (Figure S7 and Supporting file 2). We tested the impacts on dispersal directions by examining the association between environmental values and tree node locations (see methods). (Figure 6 and Table S1).

**Figure 6.**
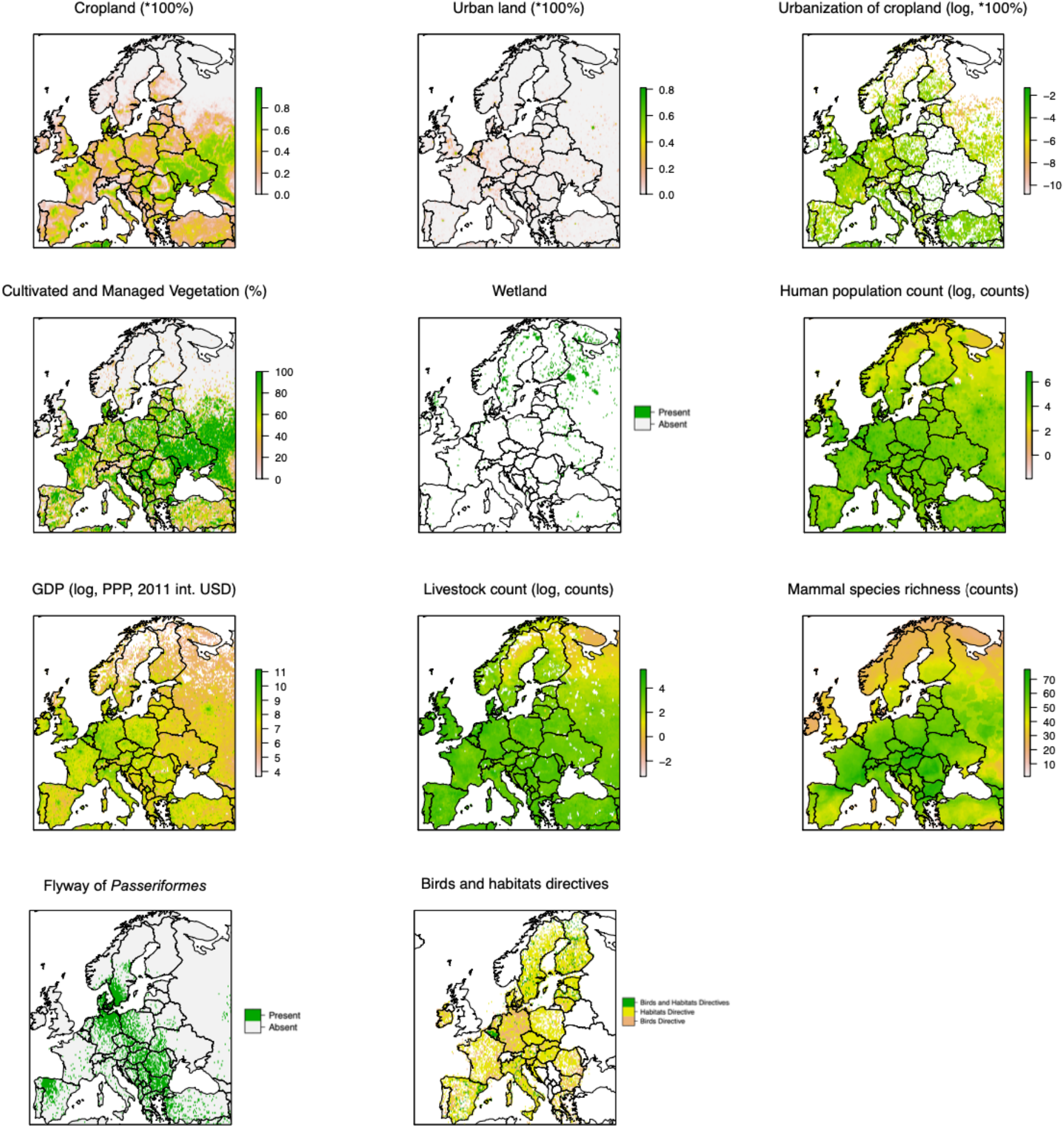
Explanatory factors significantly attracting WNV dispersal in Europe. There are eleven factors (out of the total 37 factors being tested) that may attract WNV dispersal with strong statistical support (BF>20, as shown in Table S1). Data were log-transformed where necessary for better visualization. The visualizations and full descriptions of all factors are in Figure S7 and Supporting file 2.

Our analysis found that viral lineages tended to spread towards and remained in areas with relatively high density of farming [e.g., cropland, pasture, cultivated and managed vegetation (i.e., cropland and the mixture of cropland and natural vegetation), livestock count (i.e., summed sheep, pig, horse, goat, cattle, and buffalo), and mammal richness (i.e., the number of mammal species for the entire class), Supporting file 2]. In addition, this analysis underlined that WNV lineages were more likely to spread towards areas with high urbanization status (human population density and high coverage of urban land), high coverage of wetland, as well as areas with bird directive and habitats directive and flyways of Passeriformes and Anseriformes. In addition to these impacts on directionality of spread, we found that the factors associated with high farming density (areas with high coverage of cropland, pasture, livestock count, mammal richness) may accelerate WNV dispersal velocity in Europe with strong support (BF > 20) (Figure 7 and Table S2).

**Figure 7.**
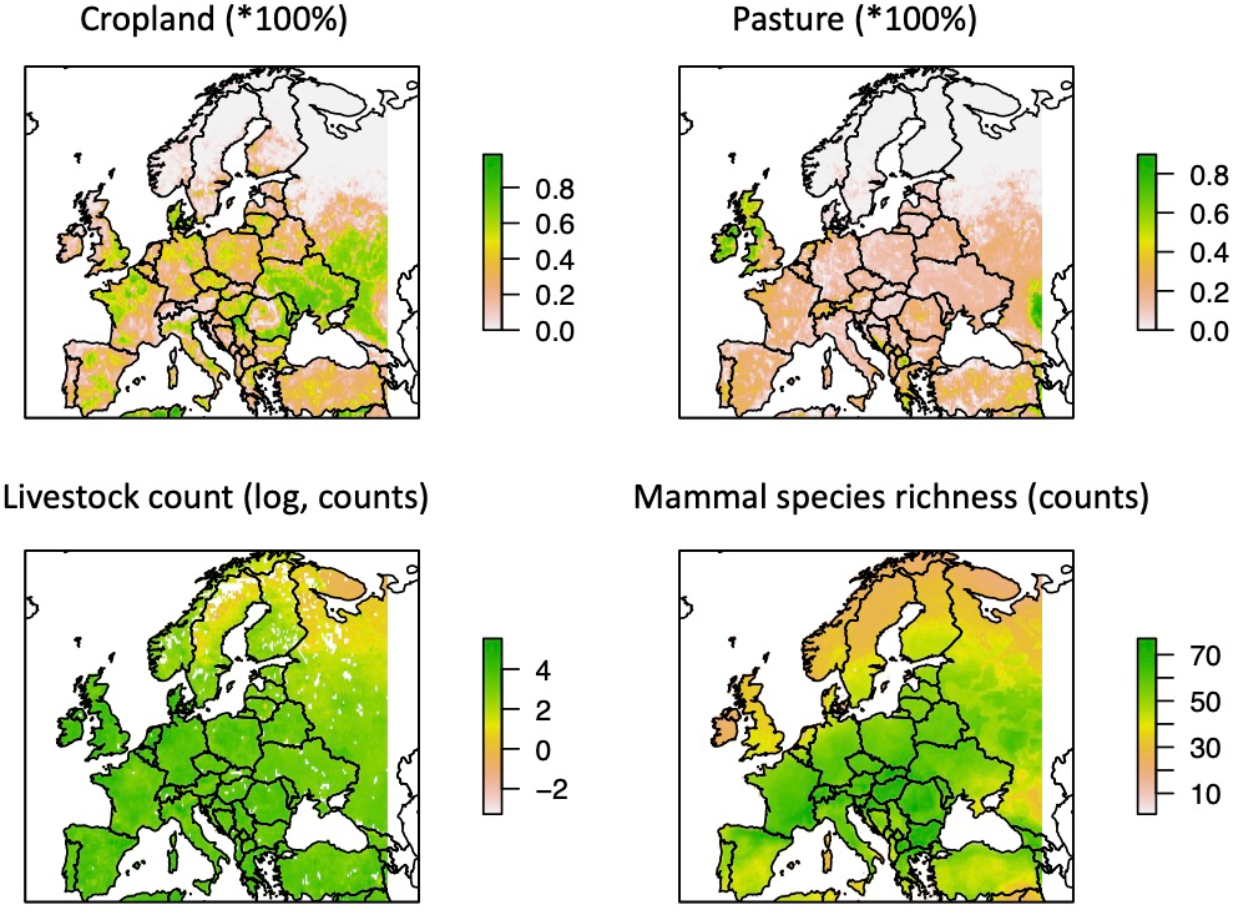
Explanatory factors have significant impact on WNV dispersal velocity in Europe. There are four factors (out of the total 37 factors being tested) that may speed up WNV dispersal with strong statistical support (BF>20, as shown in Table S2). Data were log-transformed where necessary for better visualization. The visualizations and full descriptions of all factors are in Figure S7 and Supporting file 2.

### Impact of climate/bio-diversity changes on WNV genetic diversity via time

As the dynamics of host populations (mosquitos and birds) varies widely in different months or years, it is important to understand how these variables could further affect the WNV genetic diversity or population dynamics. Therefore, we examined the relationship between WNV and climate change coefficients in a log-linear regression model. In this study, the covariates of interest included the time-varying climate related factors (including air temperature, precipitation, wind speed and direction, leaf area index) and the factors related to the trend of population changes of different types of birds in Europe (Figure S8 and Supporting file 3).

We found a significant positive association between the population history of the WNV-2a and the air temperature in Europe between 2004 and 2021. This concordance is supported by a positive, statistically significant estimate of the coefficient relating the WNV-2a effective population size to air temperature with a posterior mean of 0.18 with 95% HPD (0.08,1.68) (Figure 8 and Table S3). In contrast, wind speed in the eastward direction (rather than the northward direction) at a height of 10 and 100 meters above the surface of the Earth, monthly total precipitation, and the trends of farmland bird populations (the change in the relative abundance of 39 common bird species which are dependent on farmland for feeding and nesting and are not able to thrive in other habitats) were negatively correlated, which was in line with the observed data that mosquitoes were more abundant during the dry months compared to the wet months. In comparison, northward wind speed and the leaf area index of different types of vegetation did not have a significant impact on the genetic diversity changes of WNV-2a viruses (Table S3).

**Figure 8.**
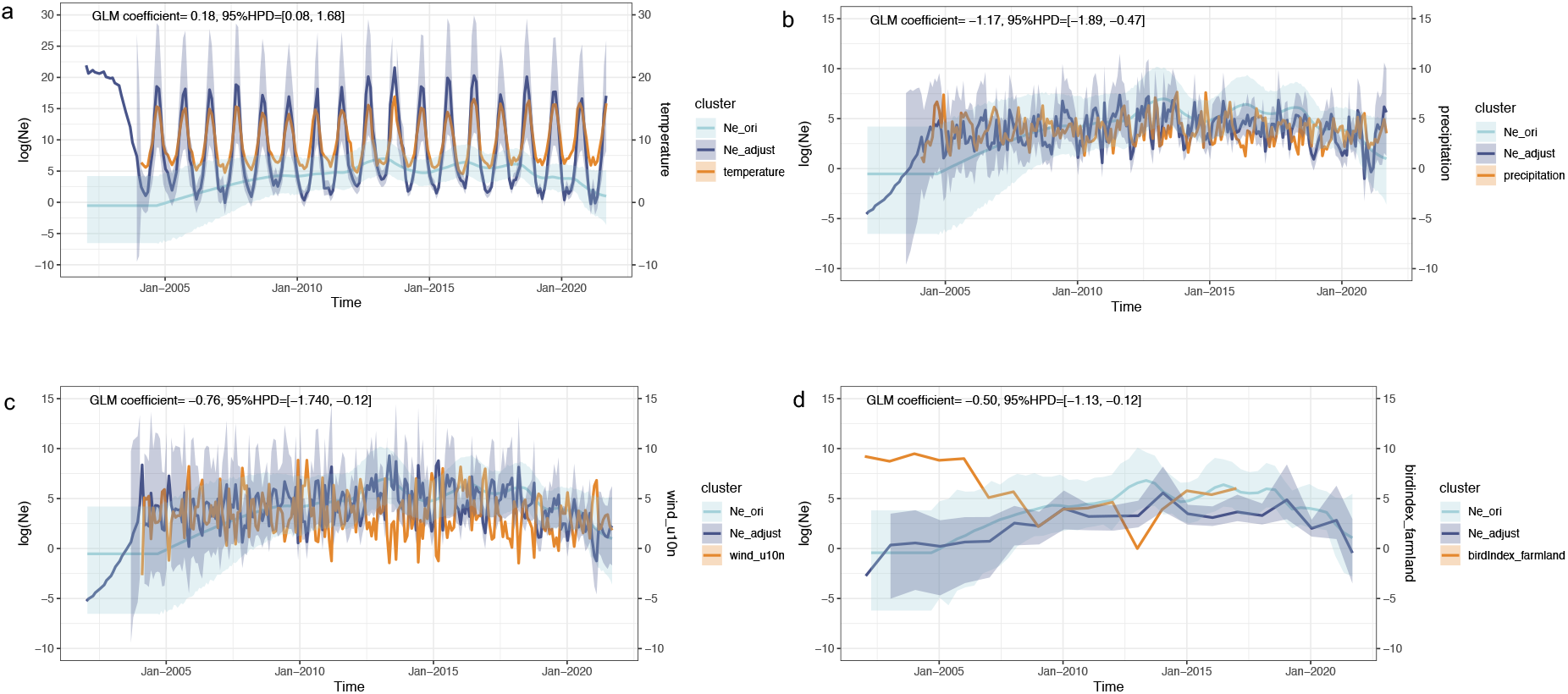
Time-varied factors with significant impacts on population dynamics via time. These associations were tested with a generalized linear model (GLM) extension of the coalescent model used to infer the dynamics of the viral effective population size of the virus (Ne) through time. We here present the following time-series variables as significant associated covariates (orange curves): a) mean temperature, b) mean precipitation, c) eastward wind speed at 10m, and d) the trend of farmland bird population index(measures the rate of change in the relative abundance of common bird species) between 2004 and 2019. Posterior mean estimates of the viral effective population size based on both sequence data and covariate data are represented by dark blue curves, and the corresponding dark blue polygon reflects the 95% HPD region. Posterior mean estimates of the viral effective population size inferred strictly from sequence data are represented by light blue curves and the corresponding light blue polygon reflects the 95% HPD region. A significant association between the covariate and effective population size is inferred when the 95% HPD interval of the GLM coefficient excludes zero. See the skygrid-GLM results of all factors being tested in Table S3. The visualizations and full descriptions of all factors being tested are in Figure S8 and Supporting file 3.

## Discussion

In this study, we used spatially explicit phylogeographic and phylodynamic inference to reconstruct the dispersal patterns and dynamics of WNV dominant in Europe in the past twenty years. Using comparative analyses of lineage dispersal statistics, we detected distinct evolutionary histories of lineages WNV-1 and WNV-2 and of sublineages WNV-2a and 2b within the overall spread of WNV. Although the evolution of WNV has previously been examined in individual European countries in different time spans ^15,24^, this study has employed a phylodynamic model to incorporate sequence data, ecological data and epidemiology data and quantified the overall evolution and dispersal history across Europe. Moreover, the presented analysis provided insight into the potential impact of predictors that influence WNV dispersal and diversity in Europe.

We described how WNV is present and spreads across Europe using viral sequence data. WNV in Europe has higher lineage diversity (six different lineages) compared to other regions in the world, e.g., only one lineage (WNV-1) has been found in North America since 1999^25,26^. The current predominant WNV-2 (2a) sequences were only found within Europe, indicating that this lineage is more likely to have emerged and circulated within Europe rather than introduced via cross-continent transmissions. However, we cannot rule out the possibilities of repeated introductions, due to under-sampling in non-European regions. The WNV-2a has evolved into 2 clusters (Cluster A and B) and spread in two directions: both originated from central Europe and then spread to the west (Cluster A) and to the south (Cluster B). In recent years, the spread patterns are becoming more complicated. Since 2018, WNV-2a (Cluster A) has unusually spread further north even up to Germany and the Netherlands.

We highlighted that areas with high levels of agricultural activities may accelerate WNV dispersal velocity as well as attract the spread direction of WNV in Europe. Meanwhile, WNV is likely to spread to areas with high status of urbanization, in line with higher abundance of *Culex pipens* (Figure S3 d), which prefers urban environments. A recent study suggested that expansion in urbanization and demography has increased the risk of infectious disease outbreaks especially in the past few decades^27^. While other native and invasive mosquito species (e.g., *Ochlerotatus caspius, Culex modestus, Aedes albopictus, Culex perexiguus)*living in natural and agricultural environments may also play an important role as WNV vectors in Europe, promoting the overwintering of WNV, especially in South and Southeast Europe ^28,29^. It is also worth noting that *Culex* hybrids from the *pipiens* and *molestus forms* have been found present in Europe, which can serve as a bridge for WNV between birds and humans, and the proportions of hybrids are differentially affected by temperature, e.g., 17% of Culex are hybrids in Greece, and the figure is 6-15% in the Netherlands^30–33^. Therefore, the different dynamic histories of WNV variants (WNV-2a and WNV-2b) might also be associated with the presence of different mosquito species, which needs further investigation. Our observations suggest that enhancing mosquito control efforts, particularly on livestock farms, may reduce the rate of WNV infections; monitoring farm operations is needed to effectively mitigate mosquito risk and WNV spread. In addition, the observation of the negative correlation between WNV population dynamics and the declines in farmland bird species in Europe in the past two decades, suggests the invasion of WNV might have an impact on the population of common bird species that are dependent on farmland for feeding and nesting. Meanwhile, the loss of habitat may force bird migration, increasing the possibility of WNV spreading to new territories.

We found a strong relationship between the presence and movement of birds and WNV spread in Europe. In terms of a natural bird reservoir, WNV sequences were obtained mainly from predatory birds (35%), songbirds (20%) and captive birds (20%) (Figure S3). WNV-2a in Europe spread at a high spread rate (88km/y to 218km/y) and, therefore, is more likely correlated with bird movement than the travel of *Culex* mosquitos (approx.500m to 2Km/y)^34,35^. We have found that WNV may be “attracted” to areas with high coverage of bird habitats and to the flyway of *Passeriformes,* particularly bird species such as the Red-backed Shrike *(Lanius collurio),* which travels at a faster rate than WNV^36–39^. Therefore, our work suggested that although the risk of WNV depends on both the presence of infected birds and the presence of mosquitoes transmitting the disease, bird movements seed the infections into mosquito populations occurring far away and introduce WNV into new regions.

In addition, climate changes in the past twenty years predicted viral genetic diversity over time. We found that higher temperature is correlated with high WNV genetic diversity during the entire history of WNV-2a spread in Europe. High temperature may stimulate the geographical expansion of suitable arbovirus regions^40^. Other epidemiological modelling studies have found that mild winters and drought have been associated with WNV disease outbreaks^40,41^. We also found that precipitation and wind speed had negative correlations, which could be explained by WNV infections mostly occurring during the mosquito season in summer^1^and also the dry season in most European countries. In addition, we found that the direction of wind also matters, as the impacts on virus genetic diversity is more likely hindered by eastward wind speed rather than northward (in Europe winds coming from the south tend to be warmer). A study also found that wind can affect mosquito host finding^42^.

Wild animals may have played a role in onward WNV transmissions, as antibodies were detected from wild animals, such as red deer, tree squirrels and wild rodents ^43,44 45^. This demonstrates that enhanced surveillance and sequencing efforts in an extended host range are needed to better understand the dispersal of WNV in more diversified environments. In addition, passive surveillance should focus on a range of wild birds especially predator birds and passerine birds. The same applies to human surveillance. Since the early 2000s, WNV circulation has been continuously monitored in some European countries with varying number of human and equine cases. Surveillance of mosquitoes and birds has proven to be useful for early detection of WNV circulation and identification of enzootic areas^17,46–49^. The available data do reflect enhanced efforts in some countries. In Greece, active mosquito surveillance, especially in the Central Macedonia Region, is being performed since 2010 when the virus was first identified in the country^17,46,50^. In Italy, a multi-species national surveillance plan was implemented since 2002, following the first WNV outbreak^13^. Arbovirus surveillance in birds has been set up in the Netherlands since 2016 with the number of screened samples increased each year (from n=952 in 2016 to n=7030 in 2021), although no WNV sequence was detected except in 2020^51^, which was mostly likely transmitted from Germany, according to our phylogeographic analysis. On the contrary, there is an insufficient amount of WNV genomes from the Central and Southeast Europe (such as Croatia^52^, Bulgaria^47^, Slovenia^53^ as well as Turkey^54,55^) as well as Southwest (Spain^56^ and Portugal^57^) where WNV have been detected in both humans and animals^2^ (Figure S1). Therefore, surveillance should include more sustained and collaborative efforts to fill in the gaps in WNV genomic sampling throughout Europe.

The risks of zoonotic diseases and their spread increase with globalisation, climate change and changes in human behaviour, giving pathogens numerous opportunities to colonise new territories and evolve into new forms. We used virus genomics to investigate the emergence of WNV in Europe, as well as its rapid spread, evolution in a new environment and establishment of endemic transmission. Most importantly, our study suggested targeted regions could deserve further investigation, including areas where WNV-2a was more likely to cluster in (with high urbanization and farming activities, high coverage of wetlands, as well as areas with the presence and movements of migrating and resident birds) and areas with higher farming density tended to accelerate the dispersal of WNV. This analysis could also be done on a more local level, and with information that we have compiled, so that we could predict where WNV would go in the future. Putting it into a larger perspective, our work could be extended to other zoonotic pathogens and guide strategies of enhanced surveillance systems for disease prevention and control.

## Methods

### Virus Isolation, Identification, and Genome Sequencing

In this study, we included unpublished WNV-2 genomes collected from Italy (n=8) and the Netherlands (n=6) between 2019 and 2021. Virus isolation, identification and sequencing are as previously described ^7,38,46^. Sequences have been submitted to GenBank with accession IDs OP561452-OP561459, OP762592-OP762597 (supporting file 1).

Apart from the new WNV genomes obtained in this study, we downloaded all available nucleotide sequences of WNV isolated from Europe as of 02 June 2022 from NCBI (www.ncbi.nlm.nih.gov). To identify any cross-continent transmission, we blast our European WNV dataset against sequences database at NCBI using Geneious Prime 2021.1.1 (https://www.geneious.com). We count in total 485 WNV sequences (from unique samples), including 226 full genomes sequences. Metadata of WNV sequences were updated and adjusted by this European collaborative consortium. For example, we have adjusted travel history of human cases (if known) to locate the original source of infection. The coordinates of samples were provided as well.

### Phylodynamic reconstructions

Sequences were aligned with MAFFT ^58^. Phylogenetic trees were first generated using IQtree ^59^ employing maximum likelihood (ML) under 1000 bootstraps. Sequences with >5% ambiguous nucleotide sites were excluded and for sequences 100% identical to one another, only one of the sequences was included. The nucleotide substitution model used for all phylogenetic analyses was HKY with a Gamma rate heterogeneity among sites with four rate categories. The temporal qualities of the sequence data were measured with TempEst v1.5 ^60^.

We further reconstructed time-scaled phylogenies of WNV-2a, which is the predominant lineage found in Europe, including all WNV-2a whole genome sequences (n=208), Ns3 gene sequences (n=275), and the Ns5 gene sequences (n=232), separately (Supporting file 1). The Ns3 gene dataset has the largest amount of sequences and covers the widest range of geographic areas (Supporting file 1 and Figure S4). Therefore, we mainly reported the results of the continuous phylogeographic analysis using Ns3 dataset in the main text. We first tested the temporal signals of the ML trees generated using WNV-2a full genomes and compared these results with analysis of the trees generated using NS3 and NS5 gene, respectively. The prior for the evolutionary clock rate of WNV was specified with a lognormal distribution and a mean of 4×10^-4^ subst/site/year with Standard Deviation of 5×10^-4^ ^25^. Phylodynamic analyses using WNV-2a sequences were conducted using time-scaled Bayesian phylogenetic methods in BEAST version 1.10.4 ^60^. The best fitted models were determined using stepping-stone sampling^61^, which resulted in the selection of HKY+Gamma+4 substitution model—an uncorrelated relaxed molecular clock model which assumes each branch has its own independent rate ^62^ and Skygrid ^63^ coalescent model. For each analysis, the Monte Carlo Markov Chain (MCMC) was run for 10^8^ steps and sampled every 10^4^ steps.

We further estimated the transmissions between countries and within countries by using linked parameters options in Beast to jointly estimate the between country transmissions using both the Ns5 gene sequences and Ns3 gene sequences (which have different phylogenies) ^60^. We used an asymmetric model and incorporated the Bayesian Stochastic Search Variable Selection procedure (BSSVS) to identify a sparse set of transmission rates that identify the statistically supported connectivity ^64^. We also estimated the expected number of transmissions (jumps) between countries and within countries using Markov rewards ^65^.

In addition, a continuous phylogenetic diffusion model with a Relaxed Random Walk extension ^19^ was further applied to explore the geographical spread of WNV in continuous space.

### Spatial and temporal drivers of WNV dispersal

We tested the associations between a set of potential predictors and the dispersal events of WNV using R package “seraphim”^20^, which has been developed to study the spatio-temporal phylogenies in an environmental context; it extracts the spatio-temporal information from a set of phylogenetic trees and uses this information to calculate and plot dispersion statistics. We prepared a list of potential predictors (n=37) which were assigned into five different groups including climatic, land cover, topography, socio-economic and biodiversity (Supporting file 2). The predictors used in this study were adapted from our previous analysis of the drivers of discovery of human infective RNA viruses^66^. We also included new drivers specifically relevant for arboviruses (e.g., wetland concentration, occurrence status of *Culex pipiens*) (Figure S7). Resolution of predictors ranged from 30” to regional level, and all data were rescaled to 0.25° where necessary. Collection of data on spatial and temporal drivers of WNV dispersal and the related modelling process were performed using the R version 3.6.3 (R Foundation for Statistical Computing, Vienna, Austria, 2020). For each of the potential predictors, we assessed the quality of the data sources used in terms of their ability to represent the underlying driver. Specifically, we assessed each driver against six characteristics using the method of Horigan et al. 2022 (paper submitted) (supporting file 2). Overall, the data sources were of very good quality, scoring well for accuracy, reliability and availability, including the factors regarding land use which the model output as influential on WNV spread. The data for *Culex pipiens* distribution was assessed to have the lowest quality amongst the factors included, due to the lack of available complete data. Future work could use instead, for example, outputs from Mosquito Alert (http://www.mosquitoalert.com/en/access-to-mosquito-alert-data-portal/), which has started collating reports of Culex sightings through citizen science since 2020. Similarly, forest-related species richness and livestock density were of lower quality; livestock density was a significant factor for WNV dispersal highlighting the need for these data sources to be regularly updated and ensured for accuracy.

Using these potential predictors, we can test the association between the dispersal speeds and directions for each branch in the phylogenetic tree, from an ancestral node to its descendants, with the equivalent paths corresponding to trajectories through the predictor maps from the ancestral node locations to the descendant locations. We tested if the virus tended to remain in areas with lower/higher environmental values, and/or the tendency of the lineages to disperse towards lower/higher values of the predictive factors, by estimating the Bayes factor (BF) comparing values explored under the inferred model with the null dispersal model simulated along the tree ^20^.

We also tested the impact on diffusion rate (or dispersal duration) by examining the association between dispersal durations and environmental distances computed for each branch of the tree. We estimated the value Q which measures the correlation between phylogenetic branch durations and environmental distances, by using the “least-cost” path model, which uses a least-cost algorithm to determine the route taken between the start and end locations of phylogenetic branch within the predictor raster cells^67^. Following the methodology described by Dellicour et al^67^, for each of the environmental factors, we generated three scaled rasters by transforming the original raster cell values with the following formula: vt = 1 + k(vo/vmax), where vt and vo are the transformed and original cell values, respectively, and vmax is the maximum cell value recorded in the raster. Here k (k = 10, 100 and 1000) is a rescaling parameter that tests different strengths of raster cells relative to the conductance (positive correlation) or resistance (negative correlation), with a minimum value set to “1”. We considered a BF value >20 as strong support for a significant correlation between the factors and dispersal durations as well as dispersal directions.

We applied the skygrid-GLM model ^27^ to jointly infer the WNV effective population size along with the coefficient that relates it to the covariate. This stands in contrast to post hoc approaches that ignore uncertainty by comparing point estimates of the effective population size and covariate values (e.g., using standard approaches for time-series comparisons). Here we examine the temporal relationship between the demographic history of WNV and the climate changes as well as the bio-diversity changes in Europe between 2004 to 2021 (Supporting file 3 and Figure S8). The temporal factors (n=22) we tested included two parts: one group was climate related data obtained from ERA5 with a spatial resolution of 0.25° (https://www.ecmwf.int/en/forecasts/datasets/reanalysis-datasets/era5) including monthly values of 2m air temperature, total precipitation, northward wind speed (100m, 10m neutral, and 10m); eastward wind speed (100m, 10m neutral, and 1), the leaf area index of high vegetation, as well as the leaf area index of low vegetation (Supporting file 3). We also obtained the common bird population index in Europe between 2004 to 2019 from The Pan-European Common Bird Monitoring Scheme (PECBMS) (https://pecbms.info/trends-and-indicators/species-trends/), and further grouped the data into 3 types (farmland, forest and others) and bird orders (extract bird species belong to 7 bird orders that matched with the bird types in our WNV sequence data:*Accipitriformes, Charadriiformes, Coliformes, Falconiformes, Galliformes, Strigiformes and Passeriformes)* independently (Supporting file 3).

## Data availability

The sequence data and metdata used in these analyses are available in supporting file 1; the predictors source data are available in supporting file 2 and 3. Raw data and R code used for the analyses are available via Figshare at link https://doi.org/10.6084/m9.figshare.21444783

## Acknowledgements

We extend a special thanks to Dr. Simon Dellicour and Dr. Mandev S. Gill from Rega Institute, for consultations of Seraphim package and skyGrid-GLM modelling.

This work is supported by European Union’s Horizon 2020 research and innovation programme under Grant No. 874735 (VEO). Barbara B. Shih was partially funded by a BBSRC Core Capability Grant BB/CCG1780/1 awarded to The Roslin Institute, and SL is additionally supported by Biotechnology and Biological Sciences Research Council (BBSRC) programme grant to Roslin Institute (Award Numbers BBS/E/D/20002173)

Greece, besides VEO: supported by the EWSMD project (code 0238/5030131) in the frame of the Operational Program Competitiveness, Entrepreneurship and Innovation 2014–2020 and by EMPROS project (code 02070). The National Reference centre for Arboviruses is supported by the National Public Health Organization in Greece.

We declare that we have no competing financial, professional, or personal interests that might have influenced the performance or presentation of the work described in this article.

## Author Contributions

LL, SL, A Papa, MPGK designed the study. MPGK and FMA designed the collaborative data mining project (VEO), convened the multidisciplinary partnership and provided critical input on study design and manuscript. LL, BBOM, EM, RSS, SP, KT, AS, SR, AG, A Pohlmann, MB, A Papa and LB set up sample and data collection and generated sequence data. LL and FZ were involved in collecting data on the spatial and temporal drivers of WNV dispersal. LL, FZ, SL, BBOM, RSS, BBS and RL were involved in data analysis and interpretation. LL, FZ, SL wrote the manuscript and MPGK, FB, FMA, BBOM, RSS, A Papa, MB, LB, MW provided critical feedback and contributed to manuscript editing. All authors gave final approval of the version to be published.

## Reference

1 Colpitts, T. M., Conway, M. J., Montgomery, R. R. & Fikrig, E. West Nile Virus: biology, transmission, and human infection. Clin Microbiol Rev 25, 635–648, doi:10.1128/CMR.00045-12 (2012).

2 Napp, S., Petric, D. & Busquets, N. West Nile virus and other mosquito-borne viruses present in Eastern Europe. Pathog Glob Health 112, 233–248, doi:10.1080/20477724.2018.1483567 (2018).

3 Kramer, L. D. in Encyclopedia of Virology (Third Edition) (eds Brian W. J. Mahy & Marc H. V. Van Regenmortel) 440–450 (Academic Press, 2008).

4 Schmidt, V. et al. Usutu virus infection in aviary birds during the cold season. Avian Pathol 50, 427–435, doi:10.1080/03079457.2021.1962003 (2021).

5 Rizzoli, A. et al. The challenge of West Nile virus in Europe: knowledge gaps and research priorities. Euro Surveill 20, doi:10.2807/1560-7917.es2015.20.20.21135 (2015).

6 Kemenesi, G. et al. Putative novel lineage of West Nile virus in Uranotaenia unguiculata mosquito, Hungary. Virusdisease 25, 500–503, doi:10.1007/s13337-014-0234-8 (2014).

7 Barzon, L. et al. Early start of seasonal transmission and co-circulation of West Nile virus lineage 2 and a newly introduced lineage 1 strain, northern Italy, June 2022. Euro Surveill 27, doi:10.2807/1560-7917.ES.2022.27.29.2200548 (2022).

8 Garcia San Miguel Rodriguez-Alarcon, L. et al. Unprecedented increase of West Nile virus neuroinvasive disease, Spain, summer 2020. Euro Surveill 26, doi:10.2807/1560-7917.ES.2021.26.19.2002010 (2021).

9 Filipe, A. R. & Pinto, M. R. Survey for antibodies to arboviruses in serum of animals from southern Portugal. Am J Trop Med Hyg 18, 423–426, do0i:10.4269/ajtmh.1969.18.423 (1969).

10 Zannoli, S. & Sambri, V. West Nile Virus and Usutu Virus Co-Circulation in Europe: Epidemiology and Implications. Microorganisms 7, doi:10.3390/microorganisms7070184 (2019).

11 Bakonyi, T. & Haussig, J. M. West Nile virus keeps on moving up in Europe. Euro Surveill 25, doi:10.2807/1560-7917.ES.2020.25.46.2001938 (2020).

12 Control, E. C. f. D. P. a. West Nile virus infection. ECDC. Annual epidemiological report for 2018 (2019).

13 Vilibic-Cavlek, T. et al. Emerging Trends in the Epidemiology of West Nile and Usutu Virus Infections in Southern Europe. Front Vet Sci 6, 437, doi:10.3389/fvets.2019.00437 (2019).

14 Lanciotti, R. S. et al. Origin of the West Nile virus responsible for an outbreak of encephalitis in the northeastern United States. Science 286, 2333–2337, doi:10.1126/science.286.5448.2333 (1999).

15 Ziegler, U. et al. West Nile Virus Epidemic in Germany Triggered by Epizootic Emergence, 2019. Viruses 12, doi:10.3390/v12040448 (2020).

16 Aguilera-Sepulveda, P. et al. West Nile Virus Lineage 2 Spreads Westwards in Europe and Overwinters in North-Eastern Spain (2017-2020). Viruses 14, doi:10.3390/v14030569 (2022).

17 Pervanidou, D. et al. West Nile virus in humans, Greece, 2018: the largest seasonal number of cases, 9 years after its emergence in the country. Euro Surveill 25, doi:10.2807/1560-7917.ES.2020.25.32.1900543 (2020).

18 Global Consortium for, H. N. & Related Influenza, V. Role for migratory wild birds in the global spread of avian influenza H5N8. Science 354, 213–217, doi:10.1126/science.aaf8852 (2016).

19 Lemey, P., Rambaut, A., Welch, J. J. & Suchard, M. A. Phylogeography takes a relaxed random walk in continuous space and time. Mol Biol Evol 27, 1877–1885, doi:10.1093/molbev/msq067 (2010).

20 Dellicour, S., Rose, R., Faria, N. R., Lemey, P. & Pybus, O. G. SERAPHIM: studying environmental rasters and phylogenetically informed movements. Bioinformatics 32, 3204–3206, doi:10.1093/bioinformatics/btw384 (2016).

21 Gill, M. S., Lemey, P., Bennett, S. N., Biek, R. & Suchard, M. A. Understanding Past Population Dynamics: Bayesian Coalescent-Based Modeling with Covariates. Syst Biol 65, 1041–1056, doi:10.1093/sysbio/syw050 (2016).

22 Fall, G. et al. Biological and phylogenetic characteristics of West African lineages of West Nile virus. PLoS Negl Trop Dis 11, e0006078, doi:10.1371/journal.pntd.0006078 (2017).

23 Papa, A. et al. Emergence of West Nile virus lineage 2 belonging to the Eastern European subclade, Greece. Arch Virol 164, 1673–1675, doi:10.1007/s00705-019-04243-8 (2019).

24 Chaintoutis, S. C., Papa, A., Pervanidou, D. & Dovas, C. I. Evolutionary dynamics of lineage 2 West Nile virus in Europe, 2004-2018: Phylogeny, selection pressure and phylogeography. Mol Phylogenet Evol 141, 106617, doi:10.1016/j.ympev.2019.106617 (2019).

25 Hadfield, J. et al. Twenty years of West Nile virus spread and evolution in the Americas visualized by Nextstrain. Plos Pathog 15, e1008042, doi:10.1371/journal.ppat.1008042 (2019).

26 Dellicour, S. et al. Epidemiological hypothesis testing using a phylogeographic and phylodynamic framework. Nat Commun 11, 5620, doi:10.1038/s41467-020-19122-z (2020).

27 Baker, R. E. et al. Infectious disease in an era of global change. Nat Rev Microbiol 20, 193–205, doi:10.1038/s41579-021-00639-z (2022).

28 Mancini, G. et al. Mosquito species involved in the circulation of West Nile and Usutu viruses in Italy. Vet Ital 53, 97–110, doi:10.12834/VetIt.114.933.4764.2 (2017).

29 Fortuna, C. et al. Evaluation of vector competence for West Nile virus in Italian Stegomyia albopicta (=Aedes albopictus) mosquitoes. Med Vet Entomol 29, 430–433, doi:10.1111/mve.12133 (2015).

30 Papa, A., Xanthopoulou, K., Tsioka, A., Kalaitzopoulou, S. & Mourelatos, S. West Nile virus in mosquitoes in Greece. Parasitol Res 112, 1551–1555, doi:10.1007/s00436-013-3302-x (2013).

31 Gomes, B. et al. Distribution and hybridization of Culex pipiens forms in Greece during the West Nile virus outbreak of 2010. Infect Genet Evol 16, 218–225, doi:10.1016/j.meegid.2013.02.006 (2013).

32 Vogels, C. B. F., Goertz, G. P., Pijlman, G. P. & Koenraadt, C. J. M. Vector competence of northern and southern European Culex pipiens pipiens mosquitoes for West Nile virus across a gradient of temperatures. Med Vet Entomol 31, 358–364, doi:10.1111/mve.12251 (2017).

33 Vogels, C. B., Fros, J. J., Goertz, G. P., Pijlman, G. P. & Koenraadt, C. J. Vector competence of northern European Culex pipiens biotypes and hybrids for West Nile virus is differentially affected by temperature. Parasit Vectors 9, 393, doi:10.1186/s13071-016-1677-0 (2016).

34 Ciota, A. T. et al. Dispersal of Culex mosquitoes (Diptera: Culicidae) from a wastewater treatment facility. J Med Entomol 49, 35–42, doi:10.1603/me11077 (2012).

35 Hamer, G. L. et al. Dispersal of adult culex mosquitoes in an urban west nile virus hotspot: a mark-capture study incorporating stable isotope enrichment of natural larval habitats. PLoS Negl Trop Dis 8, e2768, doi:10.1371/journal.pntd.0002768 (2014).

36 Ain-Najwa, M. Y. et al. Evidence of West Nile virus infection in migratory and resident wild birds in west coast of peninsular Malaysia. One Health 10, 100134, doi:10.1016/j.onehlt.2020.100134 (2020).

37 Mancuso, E. et al. West Nile and Usutu Virus Introduction via Migratory Birds: A Retrospective Analysis in Italy. Viruses 14, doi:10.3390/v14020416 (2022).

38 Rappole, J. H. & Hubalek, Z. Migratory birds and West Nile virus. J Appl Microbiol 94 Suppl, 47S–58S, doi:10.1046/j.1365-2672.94.s1.6.x (2003).

39 Lopez, G., Jimenez-Clavero, M. A., Tejedor, C. G., Soriguer, R. & Figuerola, J. Prevalence of West Nile virus neutralizing antibodies in Spain is related to the behavior of migratory birds. Vector Borne Zoonotic Dis 8, 615–621, doi:10.1089/vbz.2007.0200 (2008).

40 Di Pol, G., Crotta, M. & Taylor, R. A. Modelling the temperature suitability for the risk of West Nile Virus establishment in European Culex pipiens populations. Transbound Emerg Dis, doi:10.1111/tbed.14513 (2022).

41 Watts, M. J., Sarto, I. M. V., Mortyn, P. G. & Kotsila, P. The rise of West Nile Virus in Southern and Southeastern Europe: A spatial-temporal analysis investigating the combined effects of climate, land use and economic changes. One Health 13, 100315, doi:10.1016/j.onehlt.2021.100315 (2021).

42 Hoffmann, E. J. & Miller, J. R. Reassessment of the role and utility of wind in suppression of mosquito (Diptera: Culicidae) host finding: stimulus dilution supported over flight limitation. J Med Entomol 40, 607–614, doi:10.1603/0022-2585-40.5.607 (2003).

43 Garcia-Bocanegra, I. et al. Spatio-temporal trends and risk factors affecting West Nile virus and related flavivirus exposure in Spanish wild ruminants. BMC Vet Res 12, 249, doi:10.1186/s12917-016-0876-4 (2016).

44 Romeo, C. et al. Are tree squirrels involved in the circulation of flaviviruses in Italy? Transbound Emerg Dis 65, 1372–1376, doi:10.1111/tbed.12874 (2018).

45 Cosseddu, G. M. et al. Serological Survey of Hantavirus and Flavivirus Among Wild Rodents in Central Italy. Vector Borne Zoonotic Dis 17, 777–779, doi:10.1089/vbz.2017.2143 (2017).

46 Tsioka, K. et al. Detection and molecular characterization of West Nile virus in Culex pipiens mosquitoes in Central Macedonia, Greece, 2019-2021. Acta Trop 230, 106391, doi:10.1016/j.actatropica.2022.106391 (2022).

47 Christova, I. et al. West Nile virus lineage 2 in humans and mosquitoes in Bulgaria, 2018-2019. J Clin Virol 127, 104365, doi:10.1016/j.jcv.2020.104365 (2020).

48 Papa, A., Papadopoulou, E., Kalaitzopoulou, S., Tsioka, K. & Mourelatos, S. Detection of West Nile virus and insect-specific flavivirus RNA in Culex mosquitoes, central Macedonia, Greece. Trans R Soc Trop Med Hyg 108, 555–559, doi:10.1093/trstmh/tru100 (2014).

49 Sofia, M. et al. West Nile Virus Occurrence and Ecological Niche Modeling in Wild Bird Species and Mosquito Vectors: An Active Surveillance Program in the Peloponnese Region of Greece. Microorganisms 10, doi:10.3390/microorganisms10071328 (2022).

50 Hadjichristodoulou, C. et al. West Nile Virus Seroprevalence in the Greek Population in 2013: A Nationwide Cross-Sectional Survey. PLoS One 10, e0143803, doi:10.1371/journal.pone.0143803 (2015).

51 Sikkema, R. S. et al. Detection of West Nile virus in a common whitethroat (Curruca communis) and Culex mosquitoes in the Netherlands, 2020. Euro Surveill 25, doi:10.2807/1560-7917.ES.2020.25.40.2001704 (2020).

52 Vilibic-Cavlek, T. et al. Emerging Trends in the West Nile Virus Epidemiology in Croatia in the ‘One Health’ Context, 2011-2020. Trop Med Infect Dis 6, doi:10.3390/tropicalmed6030140 (2021).

53 Knap, N. et al. West Nile Virus in Slovenia. Viruses 12, doi:10.3390/v12070720 (2020).

54 Ergunay, K., Bakonyi, T., Nowotny, N. & Ozkul, A. Close Relationship between West Nile Virus from Turkey and Lineage 1 Strain from Central African Republic. Emerging Infectious Diseases 21, 352–355, doi:10.3201/eid2102.141135 (2015).

55 Erdogan Bamac, O. et al. Emergence of West Nile Virus Lineage-2 in Resident Corvids in Istanbul, Turkey. Vector Borne Zoonotic Dis 21, 892–899, doi:10.1089/vbz.2021.0010 (2021).

56 Figuerola, J. et al. A One Health view of the West Nile virus outbreak in Andalusia (Spain) in 2020. Emerg Microbes Infect 11, 2570–2578, doi:10.1080/22221751.2022.2134055 (2022).

57 Lourenco, J. et al. West Nile virus transmission potential in Portugal. Commun Biol 5, 6, doi:10.1038/s42003-021-02969-3 (2022).

58 Katoh, K. & Standley, D. M. MAFFT: iterative refinement and additional methods. Methods Mol Biol 1079, 131–146, doi:10.1007/978-1-62703-646-7_8 (2014).

59 Minh, B. Q. et al. IQ-TREE 2: New Models and Efficient Methods for Phylogenetic Inference in the Genomic Era. Mol Biol Evol 37, 1530–1534, doi:10.1093/molbev/msaa015 (2020).

60 Rambaut, A., Lam, T. T., Carvalho, L. M. & Pybus, O. G. Exploring the temporal structure of heterochronous sequences using TempEst (formerly Path-O-Gen). Virus Evol 2, doi:ARTN vew00710.1093/ve/vew007 (2016).

61 Baele, G. et al. Improving the Accuracy of Demographic and Molecular Clock Model Comparison While Accommodating Phylogenetic Uncertainty. Mol Biol Evol 29, 2157–2167, doi:10.1093/molbev/mss084 (2012).

62 Drummond, A. J., Ho, S. Y., Phillips, M. J. & Rambaut, A. Relaxed phylogenetics and dating with confidence. PLoS Biol 4, e88, doi:10.1371/journal.pbio.0040088 (2006).

63 Hill, V. & Baele, G. Bayesian estimation of past population dynamics in BEAST 1.10 using the Skygrid coalescent model. Mol Biol Evol, doi:10.1093/molbev/msz172 (2019).

64 Lemey, P., Rambaut, A., Drummond, A. J. & Suchard, M. A. Bayesian phylogeography finds its roots. PLoS Comput Biol 5, e1000520, doi:10.1371/journal.pcbi.1000520 (2009).

65 O’Brien, J. D., Minin, V. N. & Suchard, M. A. Learning to count: robust estimates for labeled distances between molecular sequences. Mol Biol Evol 26, 801–814, doi:msp003 [pii]10.1093/molbev/msp003 (2009).

66 Zhang, F. et al. Global discovery of human-infective RNA viruses: A modelling analysis. Plos Pathog 16, e1009079, doi:10.1371/journal.ppat.1009079 (2020).

67 Dellicour, S., Vrancken, B., Trovao, N. S., Fargette, D. & Lemey, P. On the importance of negative controls in viral landscape phylogeography. Virus Evol 4, vey023, doi:10.1093/ve/vey023 (2018).

