## Supplementary figures and tables for "West Nile Virus spread in Europe - phylogeographic pattern analysis and key drivers"

### Supplementary Tables

Table S1 Bayes factor (BF>20 in red suggest significant) supports for the association between value of potential factors (n=37) and tree node locations (remain in, leave from) as well as dispersal durations and environmental distances computed for each branch (resistance, conductance)

|  |  | Ns3_all | | Ns5_all | | ClusterA | | ClusterB | |
| --- | --- | --- | --- | --- | --- | --- | --- | --- | --- |
| Group | Risk.Factor | Remain_in | Leave_from | Remain_in | Leave_from | Remain_in | Leave_from | Remain_in | Leave_from |
| Climatic | Annual mean temperature | 0.3 | 0.3 | 0.4 | 0.7 | 0.2 | 0.1 | 1.4 | 1.6 |
| Climatic | Annual total precipitation | 0.1 | 0 | 0.1 | 0.4 | 0 | 0 | 0.6 | 0 |
| Land cover | Deciduous Broadleaf Trees | 0.2 | 0.1 | 0.2 | 0.2 | 0 | 0.8 | 1.9 | 0.2 |
| Land cover | Evergreen Broadleaf Trees | 0 | 0.2 | 0 | 0 | 0 | 0 | 0 | 0 |
| Land cover | Mixed/Other Trees | 0.4 | 0.2 | 0.9 | 0.3 | 0.1 | 0.9 | 0.1 | 0.6 |
| Land cover | Shrubs | 1.9 | 0.3 | 2.1 | 0.4 | 0.9 | 1 | 1.4 | 0.4 |
| Land cover | Evergreen/Deciduous Needleleaf Trees | 1 | 0.2 | 3 | 0.1 | 0.9 | 0.6 | 0.2 | 0.8 |
| Land cover | Cropland | >100 | 1 | 19 | 2.4 | >100 | 0.4 | >100 | 6.1 |
| Land cover | Urbanization of cropland | >100 | 1.6 | >100 | 0.8 | >100 | 2.4 | >100 | 3.3 |
| Land cover | Urbanization of pasture | 11.5 | 0.8 | 3.2 | 0.6 | 1.3 | 0.7 | >100 | 2.2 |
| Land cover | Urbanization of secondary land | 19 | 0.6 | 13.3 | 0.4 | 5.2 | 1.5 | 6.1 | 2.6 |
| Land cover | Urbanization of primary land | 0 | 0 | 0 | 0 | 0 | 0 | 0 | 0 |
| Land cover | Pasture | 2.4 | 0 | 1.9 | 0 | 0.1 | 0 | >100 | 0.1 |
| Land cover | Primary land | 0.7 | 0.6 | 0.9 | 1.4 | 0.3 | 1.5 | 0.8 | 0.2 |
| Land cover | Secondary land | 0.7 | 0 | 0.9 | 0 | 0.2 | 0.1 | 0.6 | 0.3 |
| Land cover | Urban land | >100 | >100 | >100 | >100 | >100 | >100 | >100 | >100 |
| Land cover | Cultivated and Managed Vegetation | 99 | 0.5 | 5.2 | 1.5 | 32.3 | 0.4 | 32.3 | 1.2 |
| Land cover | Regularly Flooded Vegetation | 0.4 | 0.5 | 0.3 | 0.4 | 0.4 | 0.4 | 0.3 | 0.5 |
| Land cover | Herbaceous Vegetation | 0.8 | 0.3 | 1.6 | 0.1 | 0.2 | 1.4 | 0.1 | 1.1 |
| Land cover | Open water | 0 | 0.4 | 0.2 | 0.9 | 0 | 0.8 | 0.1 | 0.4 |
| Land cover | Lake_river_reservoir | 0.6 | 0.2 | 0.5 | 0.3 | 0.3 | 0.6 | 0.4 | 0.5 |
| Land cover | Wetland_combine | 99 | 0.2 | >100 | 0.2 | >100 | 0.2 | 0.8 | 0.4 |
| Land cover | Wetland_other | >100 | 0.2 | >100 | 0.2 | >100 | 0.2 | 1 | 0.4 |
| Land cover | Wetland concentration | 2.1 | 1.0 | 9.0 | 3.2 | 6.7 | 1.2 | 0.9 | 1.6 |
| Topography | Elevation | 0 | 0 | 0.1 | 0 | 0 | 0 | 0.2 | 0 |
| Socio-economic | GDP | >100 | >100 | >100 | >100 | >100 | >100 | >100 | >100 |
| Socio-economic | Human population | >100 | >100 | >100 | >100 | >100 | >100 | >100 | >100 |
| Biodiversity | Livestock count | 24 | 1.1 | 32.3 | 0.8 | 6.7 | 0.4 | >100 | 9 |
| Biodiversity | Mammal species richness | 99 | 0.5 | 4 | 0.4 | 5.7 | 0.3 | 11.5 | 1.4 |
| Biodiversity | Flyway_Anseriformes | 14 | 0.3 | 12 | 0.1 | 10.1 | 0.6 | 0 | 0.1 |
| Biodiversity | Flyway_Apodiformes | 1 | 0.5 | 1.8 | 0.4 | 1.9 | 0.4 | 0.3 | 1.1 |
| Biodiversity | Flyway_Passeriformes | 49 | 1.3 | 3 | 2.2 | 13.3 | 6.1 | 6.7 | 0.2 |
| Biodiversity | Birds Directive | >100 | 1.9 | 24 | 1.7 | 15.7 | 2.1 | >100 | 2.8 |
| Biodiversity | Birds and Habitats Directives | 2.4 | 0.3 | 1.9 | 0.3 | 2.3 | 0.4 | 0.4 | 0.8 |
| Biodiversity | Habitats Directive | >100 | 1.6 | 49 | 2.4 | 15.7 | 2.4 | >100 | 2.8 |
| Biodiversity | Richness of forest-related species and habitats | 0.3 | 0.5 | 0.3 | 1 | 1.1 | 1 | 0 | 0.2 |
| Biodiversity | Culex pipiens status | 0 | 0 | 0 | 0 | 0 | 0 | 0 | 0 |

Table S2 Bayes factor (BF>20 in red suggests significant) supports for the association between dispersal durations and environmental distances (n=37) computed for each branch (resistance, conductance) in different scales (k=10,100,1000)

|  |  |  | Ns3 | | Ns5 | | Cluster A | | Cluster B | |
| --- | --- | --- | --- | --- | --- | --- | --- | --- | --- | --- |
| Group | Risk.Factor | k | Conductance | Resistance | Conductance | Resistance | Conductance | Resistance | Conductance | Resistance |
| Climatic | Anunal mean temperature | 10 | 0.6 | 2.3 | 2.8 | 2.3 | 0.8 | 2.1 | 1.2 | 1.1 |
| Climatic | Anunal mean temperature | 100 | 0.5 | 1.9 | 2.2 | 1.9 | 0.9 | 2.7 | 1.2 | 1.0 |
| Climatic | Anunal mean temperature | 1000 | 1.1 | 1.4 | 2.0 | 2.6 | 1.1 | 1.8 | 1.4 | 1.9 |
| Climatic | Annual total precipitation | 10 | 1.3 | 1.3 | 1.4 | 1.3 | 0.3 | 1.8 | 0.7 | 1.5 |
| Climatic | Annual total precipitation | 100 | 1.4 | 1.0 | 2.8 | 1.5 | 0.7 | 2.3 | 1.4 | 2.8 |
| Climatic | Annual total precipitation | 1000 | 1.4 | 1.0 | 1.9 | 1.5 | 0.9 | 3.8 | 1.8 | 3.3 |
| Land cover | Deciduous Broadleaf Trees | 10 | 0.9 | 3.5 | 1.9 | 19.0 | 0.3 | 8.1 | 0.8 | 1.6 |
| Land cover | Deciduous Broadleaf Trees | 100 | 0.7 | 3.5 | 6.7 | 5.7 | 0.5 | 13.3 | 0.7 | 1.9 |
| Land cover | Deciduous Broadleaf Trees | 1000 | 0.3 | 3.2 | 0.7 | 4.9 | 0.4 | 8.1 | 0.7 | 1.5 |
| Land cover | Evergreen Broadleaf Trees | 10 | 2.4 | 2.7 | 1.8 | 1.8 | 0.6 | 0.4 | 0.7 | 0.9 |
| Land cover | Evergreen Broadleaf Trees | 100 | 2.2 | 3.3 | 1.5 | 1.6 | 0.6 | 0.7 | 0.9 | 0.8 |
| Land cover | Evergreen Broadleaf Trees | 1000 | 2.4 | 3.8 | 1.3 | 2.1 | 0.5 | 0.7 | 1.3 | 0.9 |
| Land cover | Mixed/Other Trees | 10 | 3.3 | 1.4 | 0.4 | 0.9 | 1.3 | 2.6 | 1.1 | 1.3 |
| Land cover | Mixed/Other Trees | 100 | 3.5 | 1.7 | 0.3 | 1.3 | 0.5 | 2.6 | 2.1 | 1.4 |
| Land cover | Mixed/Other Trees | 1000 | 0.7 | 1.1 | 0.1 | 2.7 | 0.3 | 2.2 | 0.7 | 1.2 |
| Land cover | Shrubs | 10 | 1.4 | 0.8 | 9.0 | 1.3 | 0.6 | 0.7 | 1.9 | 0.8 |
| Land cover | Shrubs | 100 | 0.5 | 0.7 | 2.0 | 0.6 | 0.8 | 0.6 | 1.1 | 1.7 |
| Land cover | Shrubs | 1000 | 0.8 | 0.5 | 1.8 | 0.9 | 0.9 | 0.8 | 1.5 | 0.7 |
| Land cover | Evergreen/Deciduous Needleleaf Trees | 10 | 3.2 | 0.4 | 0.4 | 0.6 | 4.3 | 0.9 | 0.9 | 3.2 |
| Land cover | Evergreen/Deciduous Needleleaf Trees | 100 | 2.8 | 0.5 | 0.2 | 0.6 | 2.4 | 0.5 | 1.1 | 2.6 |
| Land cover | Evergreen/Deciduous Needleleaf Trees | 1000 | 2.8 | 0.7 | 0.3 | 1.9 | 2.4 | 0.8 | 1.1 | 2.6 |
| Land cover | Cropland | 10 | 5.3 | 1.7 | 3.5 | 4.3 | 5.7 | 0.8 | 1.4 | 1.9 |
| Land cover | Cropland | 100 | 4.9 | 1.4 | 32.3 | 1.5 | 5.3 | 1.0 | 1.9 | 1.2 |
| Land cover | Cropland | 1000 | 1.0 | 1.0 | 3.0 | 1.6 | 1.2 | 1.3 | 1.0 | 1.0 |
| Land cover | Urbanization of cropland | 10 | 1.9 | 2.3 | 2.1 | 1.6 | 1.0 | 0.8 | 0.9 | 1.0 |
| Land cover | Urbanization of cropland | 100 | 2.2 | 0.6 | 3.2 | 0.2 | 2.4 | 1.3 | 1.0 | 1.7 |
| Land cover | Urbanization of cropland | 1000 | 2.4 | 0.4 | 2.6 | 0.2 | 1.9 | 1.3 | 1.8 | 1.6 |
| Land cover | Urbanization of pasture | 10 | 1.8 | 2.7 | 2.7 | 2.0 | 0.6 | 1.3 | 0.9 | 1.2 |
| Land cover | Urbanization of pasture | 100 | 6.1 | 1.9 | 5.7 | 1.1 | 0.8 | 1.7 | 0.8 | 1.1 |
| Land cover | Urbanization of pasture | 1000 | 5.7 | 0.9 | 8.1 | 0.9 | 1.5 | 1.1 | 2.1 | 1.0 |
| Land cover | Urbanization of secondary land | 10 | 1.9 | 2.0 | 2.1 | 2.2 | 0.6 | 1.4 | 0.9 | 0.9 |
| Land cover | Urbanization of secondary land | 100 | 1.4 | 1.3 | 2.2 | 1.6 | 1.9 | 0.8 | 0.7 | 1.3 |
| Land cover | Urbanization of secondary land | 1000 | 0.9 | 1.2 | 0.4 | 0.8 | 1.8 | 0.8 | 0.6 | 0.9 |
| Land cover | Urbanization of primary land | 10 | 4.3 | 2.1 | 3.0 | 1.8 | 0.6 | 0.9 | 1.2 | 0.9 |
| Land cover | Urbanization of primary land | 100 | 4.3 | 2.7 | 4.6 | 2.1 | 0.5 | 0.9 | 1.8 | 1.0 |
| Land cover | Urbanization of primary land | 1000 | 2.8 | 2.4 | 5.7 | 1.4 | 0.5 | 1.9 | 1.3 | 0.9 |
| Land cover | Pasture | 10 | 8.1 | 0.9 | 2.0 | 3.0 | 2.3 | 1.4 | 1.7 | 2.1 |
| Land cover | Pasture | 100 | 32.3 | 1.2 | 9.0 | 1.6 | 3.2 | 2.1 | 9.0 | 1.8 |
| Land cover | Pasture | 1000 | 1.8 | 0.9 | 3.2 | 1.4 | 1.7 | 2.2 | 1.8 | 0.9 |
| Land cover | Primary land | 10 | 1.4 | 1.7 | 1.9 | 0.9 | 0.4 | 2.8 | 4.6 | 3.8 |
| Land cover | Primary land | 100 | 1.1 | 1.9 | 1.5 | 0.5 | 0.2 | 3.5 | 3.5 | 2.3 |
| Land cover | Primary land | 1000 | 1.3 | 0.4 | 2.3 | 0.3 | 0.3 | 3.3 | 2.1 | 1.0 |
| Land cover | Secondary land | 10 | 1.3 | 2.8 | 0.7 | 13.3 | 4.6 | 3.5 | 0.8 | 5.3 |
| Land cover | Secondary land | 100 | 2.8 | 4.6 | 2.8 | 3.0 | 3.2 | 4.3 | 1.0 | 3.2 |
| Land cover | Secondary land | 1000 | 0.8 | 2.8 | 1.0 | 3.5 | 1.1 | 3.3 | 0.8 | 5.7 |
| Land cover | Urban land | 10 | 4.9 | 5.7 | 3.3 | 6.7 | 3.0 | 1.2 | 1.1 | 1.3 |
| Land cover | Urban land | 100 | 1.7 | 0.7 | 1.2 | 0.3 | 3.8 | 0.1 | 0.5 | 0.6 |
| Land cover | Urban land | 1000 | 0.4 | 0.2 | 0.4 | 0.3 | 9.0 | 0.1 | 0.3 | 0.1 |
| Land cover | Cultivated and Managed Vegetation | 10 | 6.1 | 1.4 | 13.3 | 3.3 | 3.0 | 1.3 | 1.5 | 1.7 |
| Land cover | Cultivated and Managed Vegetation | 100 | 1.0 | 1.3 | 3.0 | 2.3 | 1.9 | 1.5 | 0.8 | 1.5 |
| Land cover | Cultivated and Managed Vegetation | 1000 | 0.5 | 1.6 | 0.3 | 2.1 | 1.1 | 1.1 | 0.4 | 1.0 |
| Land cover | Regularly Flooded Vegetation | 10 | 1.9 | 2.8 | 1.2 | 1.9 | 1.3 | 0.5 | 0.8 | 0.9 |
| Land cover | Regularly Flooded Vegetation | 100 | 2.8 | 3.0 | 1.7 | 1.8 | 2.0 | 0.9 | 1.4 | 1.0 |
| Land cover | Regularly Flooded Vegetation | 1000 | 3.5 | 2.1 | 2.1 | 3.5 | 4.0 | 1.4 | 1.8 | 1.6 |
| Land cover | Herbaceous Vegetation | 10 | 4.3 | 1.1 | 6.7 | 0.4 | 1.3 | 1.0 | 2.0 | 2.3 |
| Land cover | Herbaceous Vegetation | 100 | 2.1 | 1.0 | 0.4 | 0.2 | 2.1 | 1.6 | 1.2 | 5.7 |
| Land cover | Herbaceous Vegetation | 1000 | 1.6 | 0.5 | 0.2 | 0.2 | 1.4 | 1.2 | 0.7 | 1.1 |
| Land cover | Open water | 10 | 6.7 | 15.7 | 5.7 | 8.1 | 1.9 | 4.0 | 1.9 | 1.6 |
| Land cover | Open water | 100 | 3.0 | 0.6 | 5.3 | 0.6 | 3.0 | 1.1 | 1.4 | 1.0 |
| Land cover | Open water | 1000 | 2.3 | 0.4 | 4.3 | 0.1 | 5.3 | 0.7 | 1.3 | 0.9 |
| Land cover | Lake_river_reservoir | 10 | 0.9 | 1.2 | 1.4 | 0.5 | 1.2 | 0.6 | 1.4 | 0.9 |
| Land cover | Lake_river_reservoir | 100 | 1.2 | 0.9 | 1.5 | 0.8 | 1.6 | 1.3 | 0.8 | 0.4 |
| Land cover | Lake_river_reservoir | 1000 | 1.0 | 0.8 | 1.7 | 0.8 | 1.1 | 1.1 | 1.4 | 0.5 |
| Land cover | Wetland_combine | 10 | 0.5 | 0.6 | 0.9 | 0.2 | 0.6 | 0.3 | 0.7 | 0.6 |
| Land cover | Wetland_combine | 100 | 0.4 | 0.1 | 1.4 | 0.0 | 0.5 | 0.0 | 0.6 | 0.4 |
| Land cover | Wetland_combine | 1000 | 0.3 | 0.4 | 1.0 | 0.3 | 0.5 | 0.4 | 0.5 | 0.5 |
| Land cover | Wetland_other | 10 | 0.3 | 0.9 | 0.7 | 0.0 | 0.4 | 0.2 | 0.4 | 0.9 |
| Land cover | Wetland_other | 100 | 0.2 | 0.0 | 0.7 | 0.0 | 0.3 | 0.1 | 0.4 | 0.2 |
| Land cover | Wetland_other | 1000 | 0.4 | 0.3 | 0.6 | 0.1 | 0.3 | 0.2 | 0.3 | 0.2 |
| Land cover | Wetland concentration | 10 | 8.1 | 7.3 | 4.3 | 3.2 | 2.8 | 2.0 | 5.3 | 4.3 |
| Land cover | Wetland concentration | 100 | 6.7 | 6.1 | 4.9 | 2.7 | 1.7 | 2.8 | 4.6 | 4.0 |
| Land cover | Wetland concentration | 1000 | 3.0 | 1.8 | 3.8 | 0.6 | 1.4 | 0.5 | 5.3 | 1.8 |
| Topography | Elevation | 10 | 0.6 | 2.8 | 0.4 | 6.1 | 0.5 | 7.4 | 0.8 | 3.8 |
| Topography | Elevation | 100 | 0.8 | 4.6 | 0.1 | 1.6 | 0.3 | 6.1 | 1.3 | 3.2 |
| Topography | Elevation | 1000 | 0.7 | 4.9 | 0.1 | 2.4 | 0.5 | 5.7 | 0.8 | 2.6 |
| Socio-economic | GDP | 10 | 11.5 | 3.6 | 14.0 | 3.0 | 3.3 | 3.7 | 3.8 | 3.5 |
| Socio-economic | GDP | 100 | 7.3 | 3.0 | 13.3 | 1.6 | 6.7 | 0.9 | 4.6 | 3.3 |
| Socio-economic | GDP | 1000 | 1.7 | 0.0 | 1.0 | 0.0 | 8.1 | 0.0 | 2.2 | 0.3 |
| Socio-economic | Human population | 10 | 1.9 | 2.4 | 4.0 | 4.9 | 1.3 | 0.8 | 0.9 | 0.5 |
| Socio-economic | Human population | 100 | 1.4 | 1.9 | 2.2 | 4.0 | 3.5 | 0.1 | 0.9 | 0.4 |
| Socio-economic | Human population | 1000 | 1.3 | 0.3 | 1.1 | 0.2 | 5.7 | 0.3 | 0.9 | 0.3 |
| Biodiversity | Livestock count | 10 | 13.3 | 6.1 | 13.3 | 10.1 | 2.2 | 1.2 | 5.3 | 2.6 |
| Biodiversity | Livestock count | 100 | 5.3 | 1.9 | 24.0 | 0.5 | 2.3 | 0.1 | 10.1 | 1.1 |
| Biodiversity | Livestock count | 1000 | 4.6 | 0.3 | 32.3 | 0.1 | 3.5 | 0.1 | 8.1 | 0.3 |
| Biodiversity | Mammal species richness | 10 | 9.0 | 6.7 | 4.0 | 15.7 | 1.8 | 1.7 | 1.1 | 3.0 |
| Biodiversity | Mammal species richness | 100 | 24.0 | 8.1 | 3.5 | 13.3 | 2.6 | 1.9 | 2.8 | 2.1 |
| Biodiversity | Mammal species richness | 1000 | 13.3 | 5.3 | 2.7 | 11.5 | 2.0 | 2.3 | 3.3 | 2.7 |
| Biodiversity | Flyway_Anseriformes | 10 | 0.2 | 0.0 | 2.6 | 0.1 | 0.2 | 0.1 | 1.0 | 1.3 |
| Biodiversity | Flyway_Anseriformes | 100 | 0.1 | 0.9 | 0.9 | 0.0 | 0.0 | 0.1 | 1.1 | 2.0 |
| Biodiversity | Flyway_Anseriformes | 1000 | 0.0 | 2.7 | 0.8 | 0.1 | 0.1 | 0.4 | 0.9 | 1.7 |
| Biodiversity | Flyway_Apodiformes | 10 | 0.1 | 2.4 | 0.1 | 0.8 | 0.2 | 1.7 | 0.9 | 1.2 |
| Biodiversity | Flyway_Apodiformes | 100 | 0.1 | 1.0 | 0.0 | 0.4 | 0.3 | 1.1 | 1.3 | 1.6 |
| Biodiversity | Flyway_Apodiformes | 1000 | 0.1 | 1.1 | 0.0 | 0.2 | 0.3 | 1.2 | 1.4 | 0.8 |
| Biodiversity | Flyway_Passeriformes | 10 | 9.0 | 0.3 | 1.0 | 0.3 | 1.7 | 0.2 | 0.4 | 2.0 |
| Biodiversity | Flyway_Passeriformes | 100 | 3.2 | 0.2 | 2.4 | 0.4 | 0.9 | 0.2 | 0.5 | 2.1 |
| Biodiversity | Flyway_Passeriformes | 1000 | 2.4 | 0.2 | 4.0 | 0.5 | 1.0 | 1.0 | 0.4 | 1.9 |
| Biodiversity | Birds Directive | 10 | 1.6 | 0.9 | 0.1 | 0.1 | 3.0 | 1.1 | 2.7 | 2.5 |
| Biodiversity | Birds Directive | 100 | 2.1 | 0.6 | 0.1 | 0.2 | 2.4 | 0.5 | 2.6 | 2.6 |
| Biodiversity | Birds Directive | 1000 | 2.3 | 0.6 | 0.1 | 0.3 | 3.2 | 0.5 | 2.4 | 2.6 |
| Biodiversity | Birds and Habitats Directives | 10 | 0.8 | 1.1 | 0.1 | 0.1 | 3.0 | 1.4 | 2.1 | 1.4 |
| Biodiversity | Birds and Habitats Directives | 100 | 0.8 | 1.0 | 0.0 | 0.5 | 1.6 | 0.6 | 2.2 | 0.6 |
| Biodiversity | Birds and Habitats Directives | 1000 | 0.7 | 2.6 | 0.1 | 2.8 | 1.7 | 1.9 | 2.6 | 1.9 |
| Biodiversity | Habitats Directive | 10 | 1.6 | 0.8 | 0.1 | 0.1 | 3.8 | 1.2 | 2.8 | 2.7 |
| Biodiversity | Habitats Directive | 100 | 2.0 | 0.6 | 0.1 | 0.2 | 4.0 | 0.5 | 2.5 | 1.3 |
| Biodiversity | Habitats Directive | 1000 | 2.6 | 0.5 | 0.1 | 0.2 | 3.8 | 0.5 | 2.6 | 1.0 |
| Biodiversity | Richness of forest-related species and habitats | 10 | 4.3 | 3.3 | 2.4 | 2.7 | 1.1 | 0.5 | 2.0 | 2.4 |
| Biodiversity | Richness of forest-related species and habitats | 100 | 2.1 | 3.1 | 2.2 | 2.1 | 0.0 | 0.4 | 2.2 | 2.2 |
| Biodiversity | Richness of forest-related species and habitats | 1000 | 1.1 | 3.0 | 2.1 | 2.0 | 0.0 | 0.3 | 2.1 | 2.3 |
| Biodiversity | Culex pipiens status | 10 | 2.7 | 2.7 | 3.5 | 2.8 | 0.4 | 0.7 | 1.6 | 1.8 |
| Biodiversity | Culex pipiens status | 100 | 2.7 | 3.3 | 3.2 | 3.0 | 0.6 | 0.4 | 1.9 | 1.8 |
| Biodiversity | Culex pipiens status | 1000 | 2.8 | 2.6 | 3.3 | 2.7 | 0.7 | 0.8 | 1.9 | 1.9 |

Table S3 Association between temporal factors (n=22) and WNV genetic diversity via time (significant positive correlation in red and negative correlation in green)

| Covariates | time unit | coefficient mean | 95%HPD lower | 95%HPD upper |
| --- | --- | --- | --- | --- |
| Surface air temperature at 2m | monthly | 0.18 | 0.08 | 1.68 |
| Precipitation | monthly | -1.17 | -1.89 | -0.47 |
| Leaf area index, low vegetation | monthly | 1.22 | -0.62 | 3.11 |
| Leaf area index, high vegetation | monthly | 1.15 | -0.81 | 3.04 |
| 10m u-component of wind (eastward) | monthly | -0.70 | -1.29 | -0.07 |
| 100m u-component of wind (eastward) | monthly | -0.76 | -1.74 | -0.12 |
| 10m v-component of wind (northward) | monthly | -0.75 | -1.51 | 0.08 |
| 100m v-component of wind (northward) | monthly | -0.60 | -1.75 | 0.68 |
| 10m u-component of neutral wind (eastward) | monthly | -1.00 | -1.72 | -0.24 |
| 10m v-component of neutral wind (northward) | monthly | -0.56 | -1.65 | 0.64 |
| 10m wind speed | monthly | -1.11 | -1.85 | -0. 40 |
| Instantaneous 10m wind | monthly | 0.93 | -0.22 | 1.66 |
| birdindex-farmland birds | yearly | -0.50 | -1.13 | -0.12 |
| birdindex-forest birds | yearly | -0.16 | -2.06 | 1.83 |
| birdindex-other birds | yearly | -0.31 | -0.76 | 0.16 |
| birdindex-Galliformes | yearly | 0.00 | -0.14 | 0.13 |
| birdindex-Passeriformes | yearly | -0.29 | -0.88 | 0.25 |
| birdindex-Strigiformes | yearly | -0.04 | -0.16 | 0.08 |
| birdindex-Accipitriformes | yearly | -0.03 | -0.08 | 0.01 |
| birdindex-Galliformes | yearly | -0.03 | -0.20 | 0.19 |
| birdindex-Coliformes | yearly | -0.09 | -0.40 | 0.24 |
| birdindex-Falconiformes | yearly | -0.19 | -2.22 | 1.65 |

### Supplementary Figures

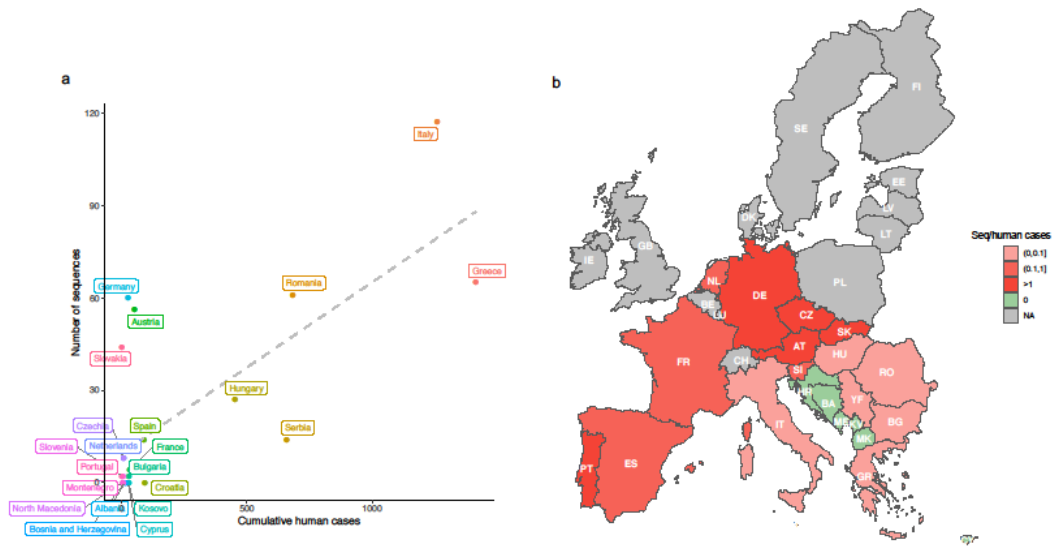

Figure S1 Surveillance and sequencing effort of WNV in Europe. (a) Comparison between cumulative human cases reported by ECDC (between 2008-2021, total n=4188) and the number of WNV sequences (between 1971-2021, total n=485) isolated from 22 different countries. (b) The sequencing effort (ratio of the number of sequences available to the number of human cases reported) per country is shown on the map: red from light to dark indicated the ratio from low to high; green indicated no sequence available although human cases have been reported; grey indicated neither human cases nor sequences are available.

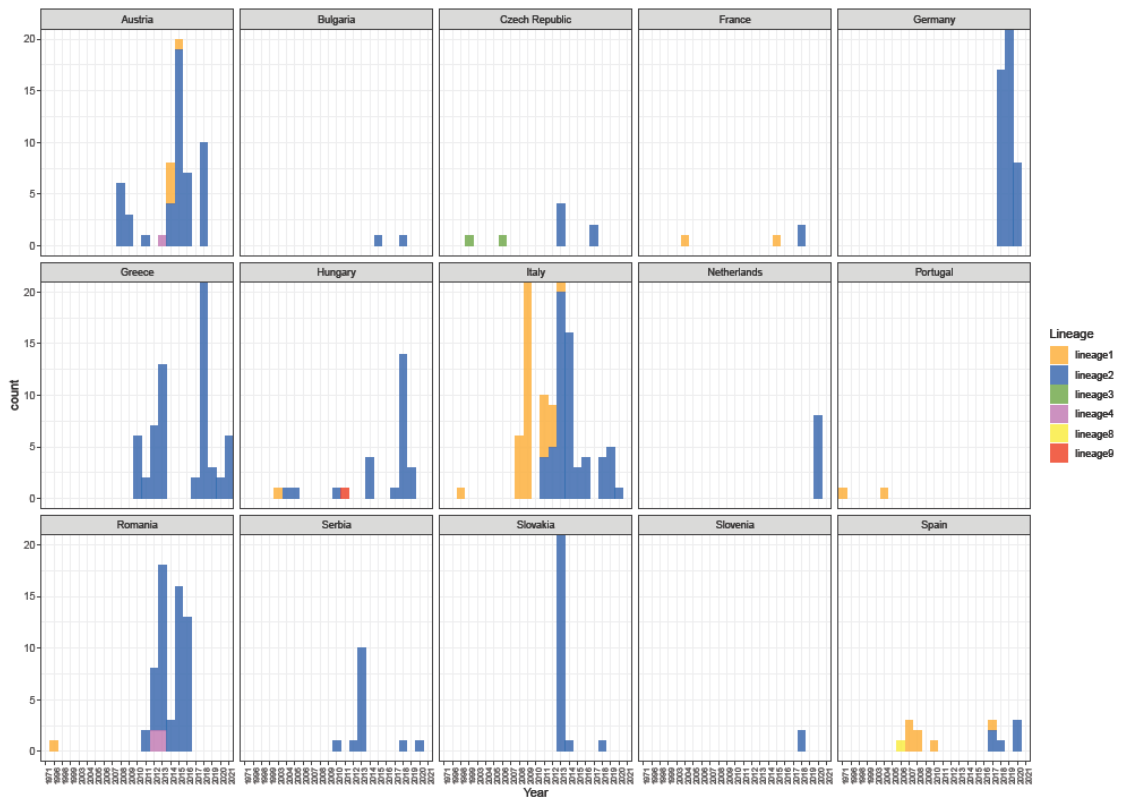

Figure S2 Number of sequences per lineage in different country and time.

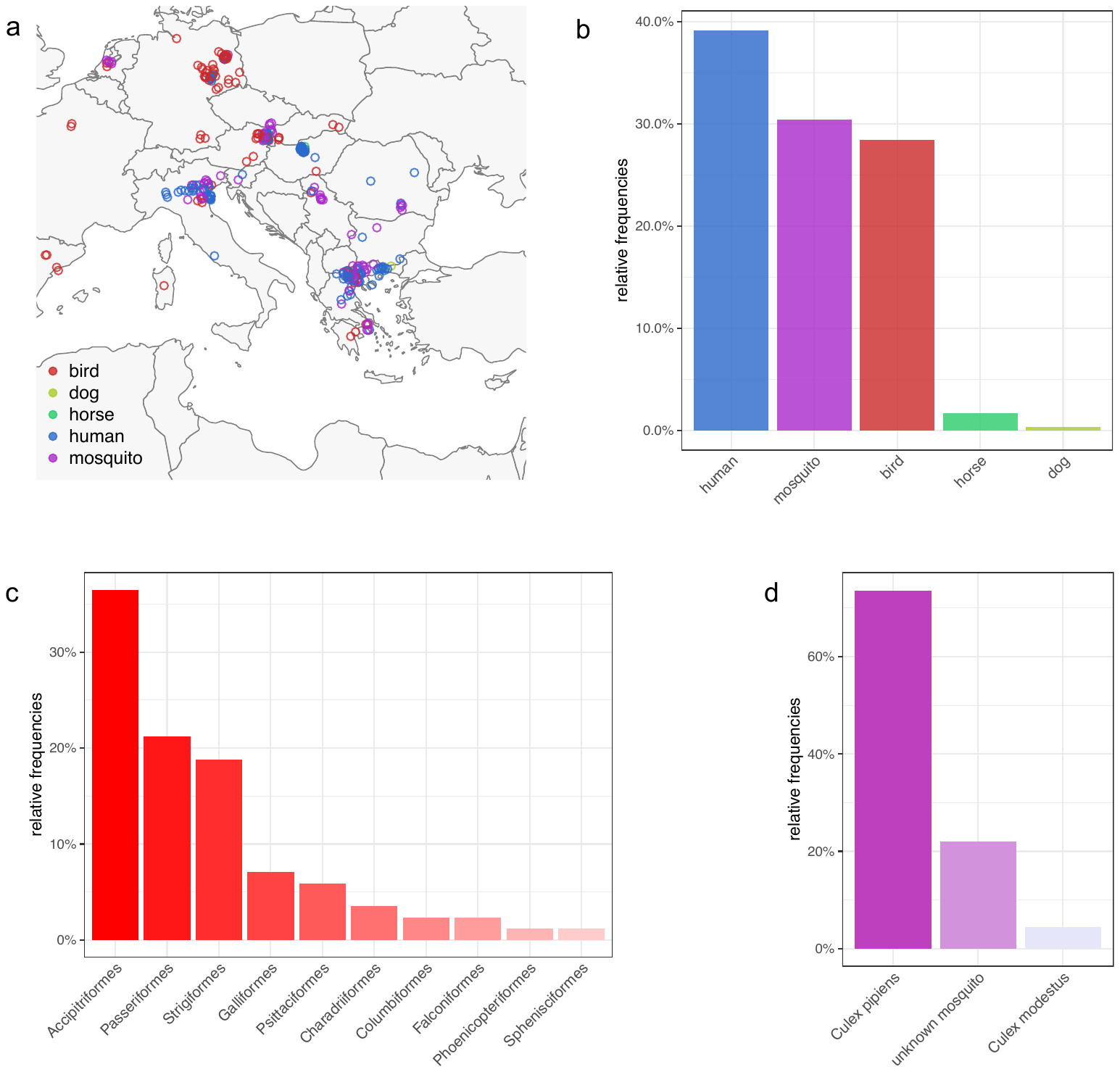
Figure S3 Distributions of WNV-2a sequences from different hosts. (a) Sequences from different hosts on maps. The colors for hosts are consistent in a-d. (b) Relative frequencies of sequences sampled from 5 different hosts. (c) Relative frequencies of sequences sampled from different bird orders. (d) Relative frequencies of sequences sampled from different mosquito species.

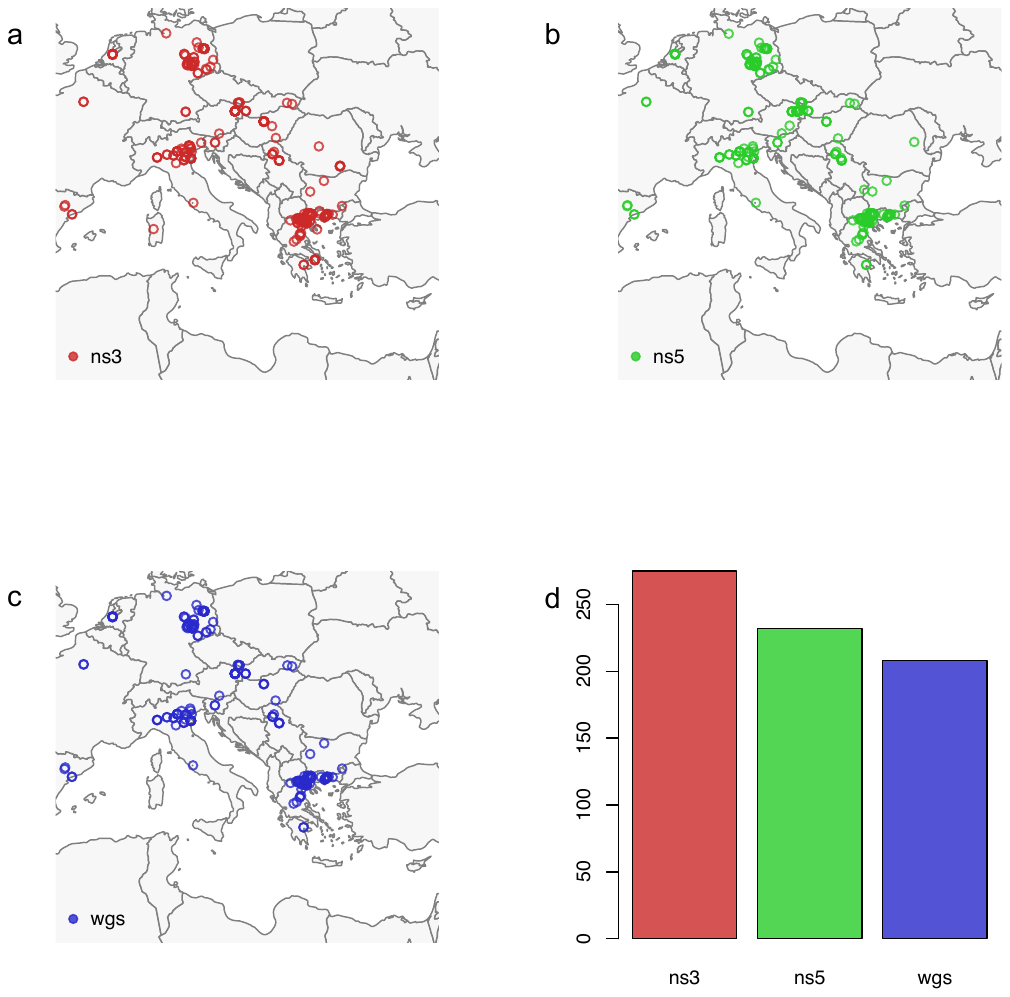

Figure S4 Geographic distributions of WNV-2a sequences dataset. a) ns3 gene, b) ns5 gene and c) full genomes and d) the number of sequences for each dataset.

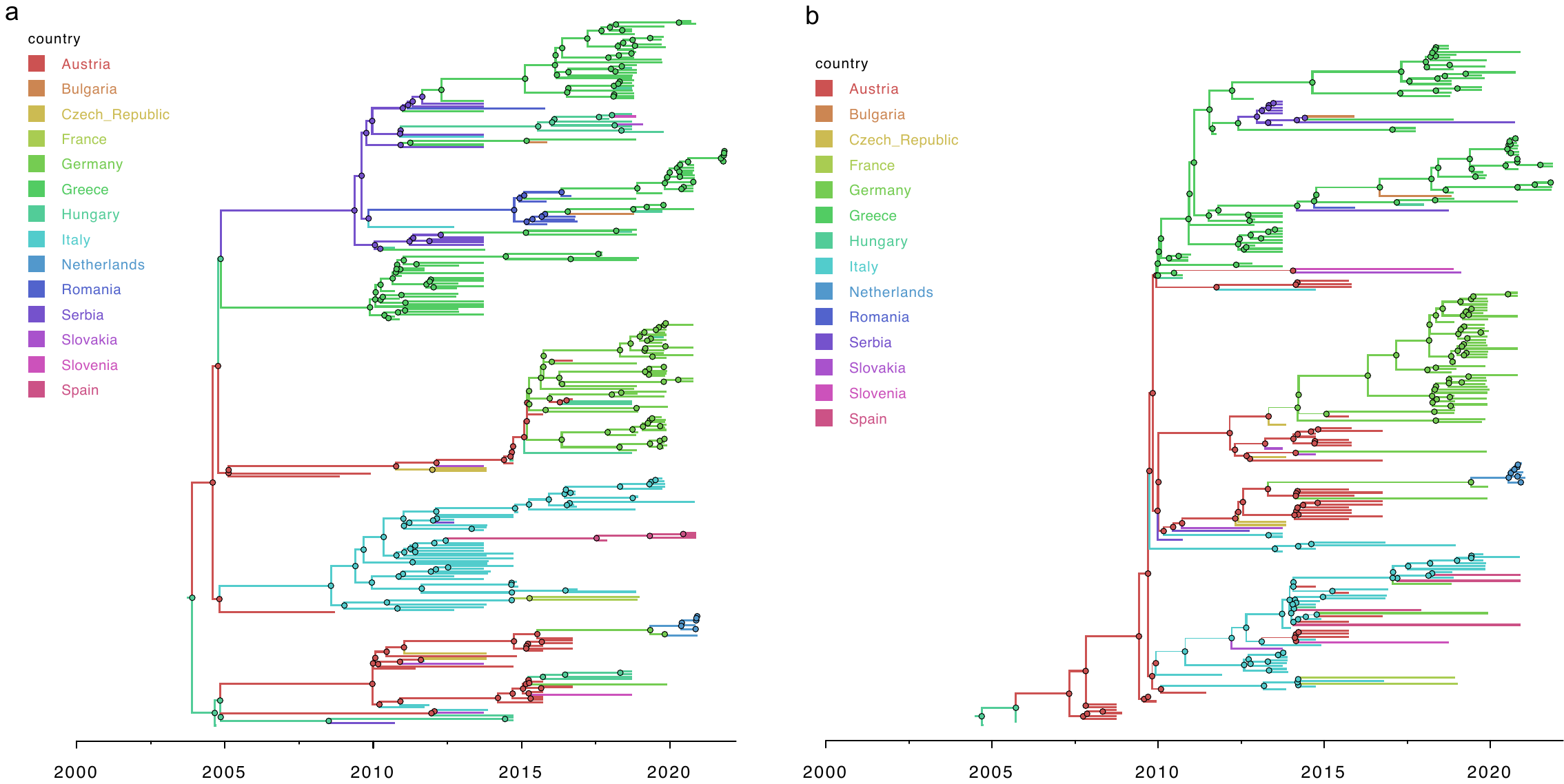

Figure S5 Time-scaled MCC tree of WNV sequences of ns3 gene (a) and ns5 gene (b) mapping with countries (n=14).

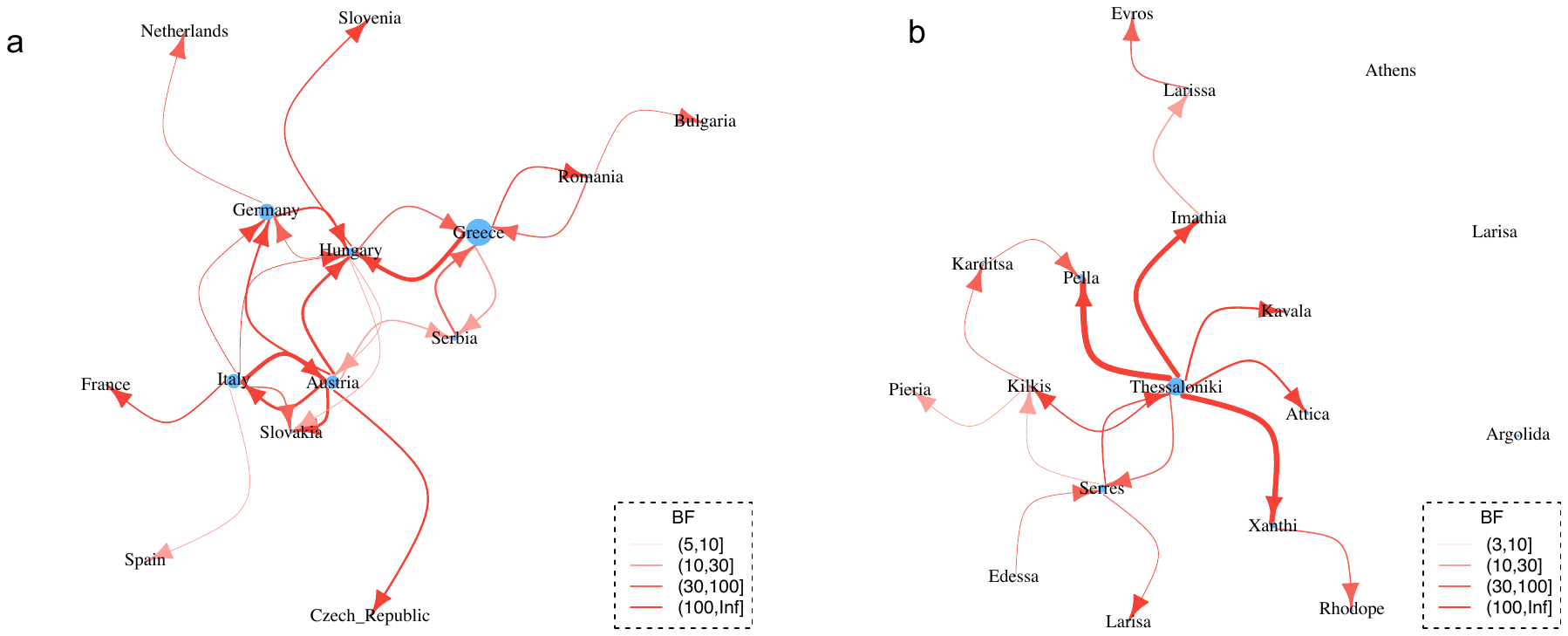
Figure S6 Transmission network inferred from the joint analysis of ns3 and ns5 phylogenies of WNV-2a sequences. (a) between European countries. (b) between regions in Greece. Size of node indicates number of samples; edge weight indicates median number of transmissions between pairs of locations; arrow on edge indicates transmission direction; color of edge from light to dark indicates Bayes Factor (BF) support from low to high only transmissions with BF >5 are shown). The correlated farms are grouped together. Nodes with no link to the others indicated no significant transmissions with other areas although sequences have been sampled.

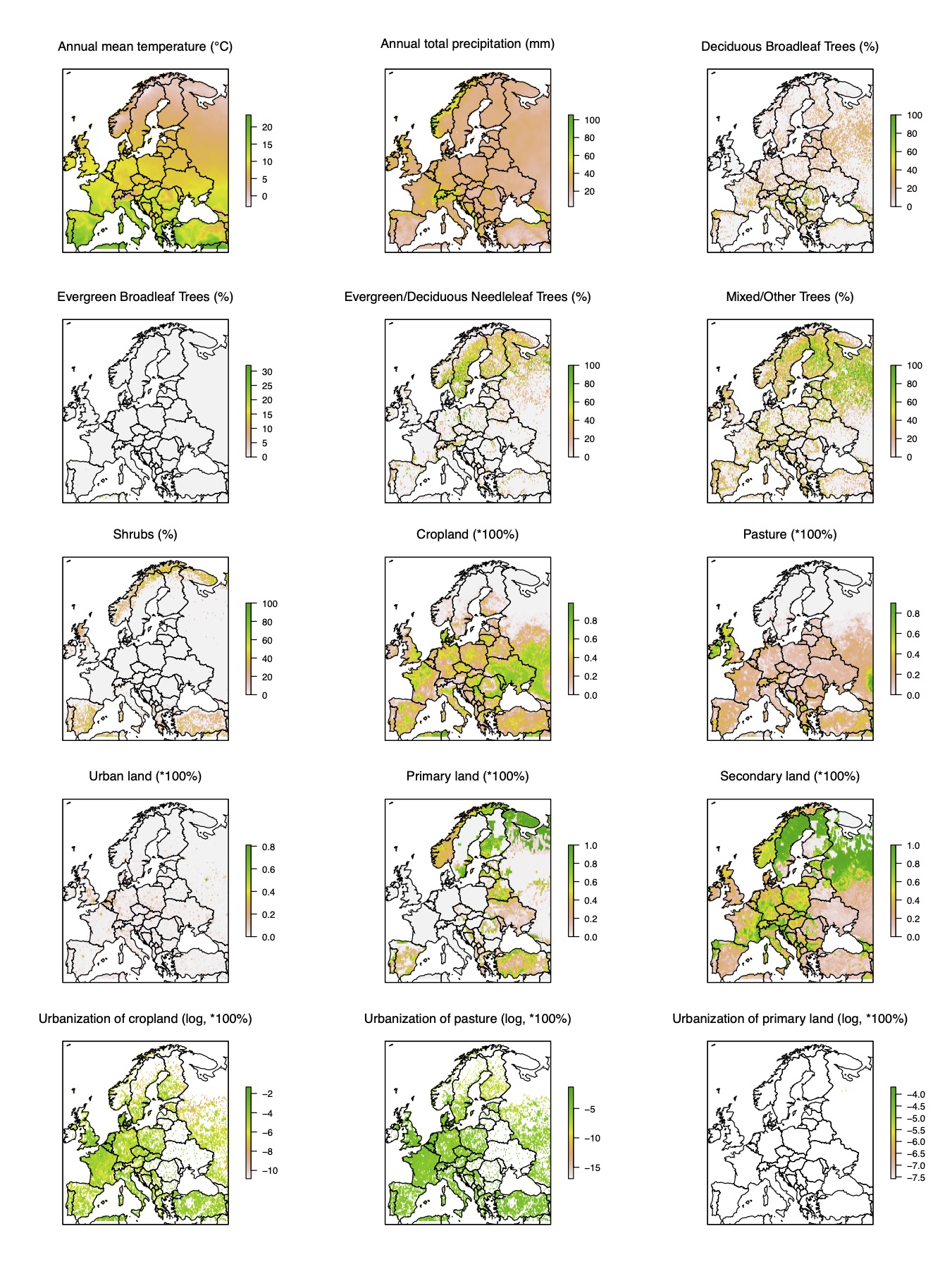

Figure S7 Distribution of predictors for dispersal of West Nile virus in Europe. Definition for each predictor is shown in Supporting file 2. Data were log-transformed where necessary for better visualization.

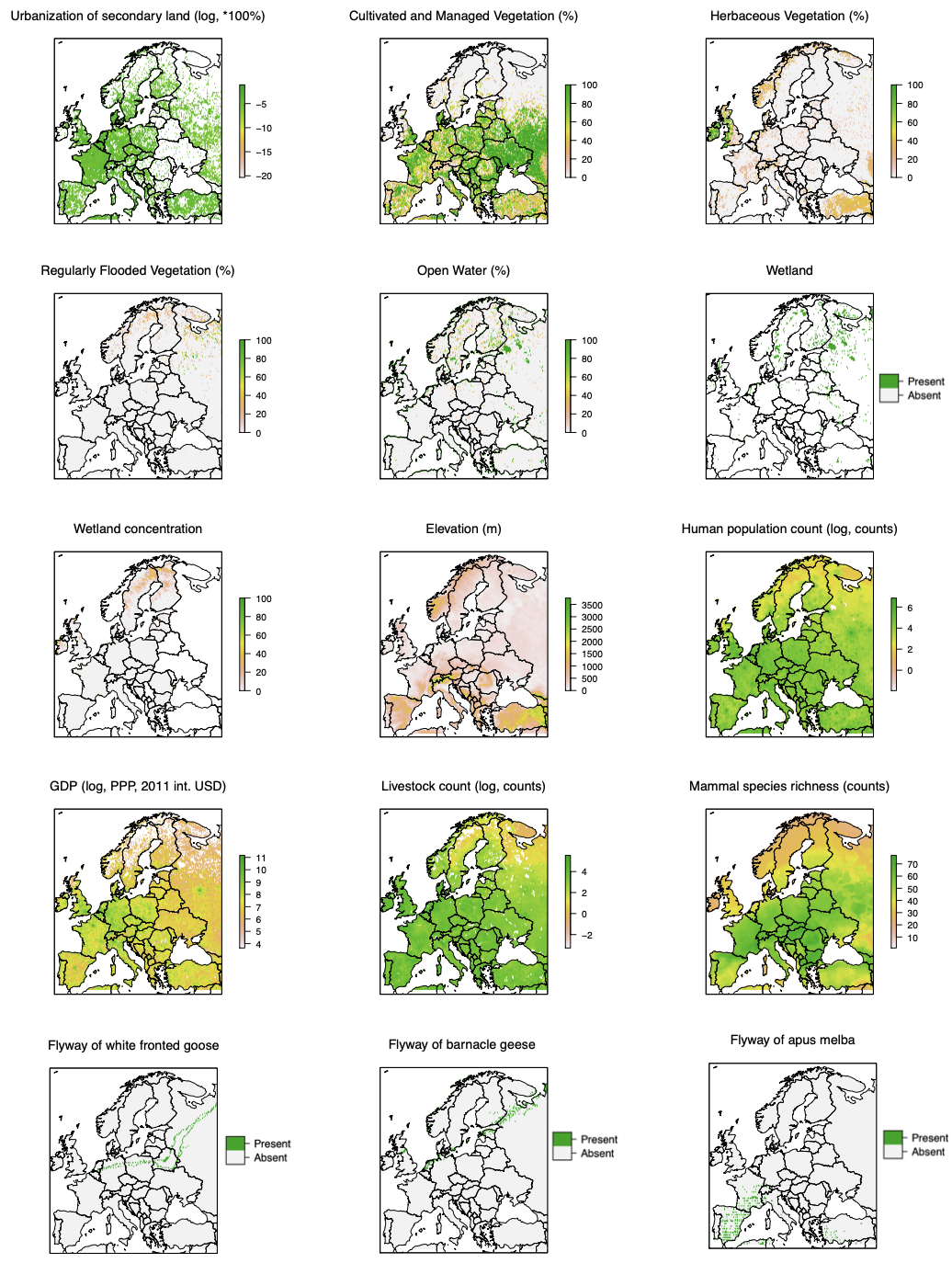

Figure S7 continued - 1

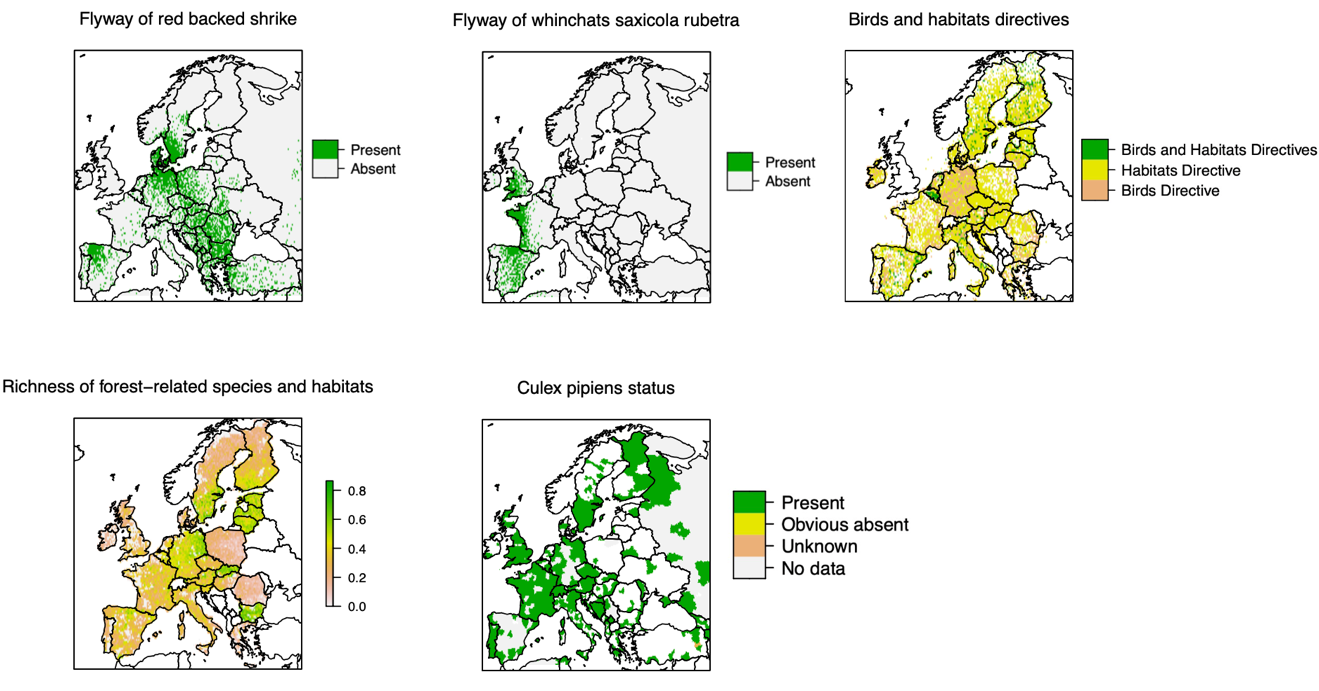

Figure S7 continued - 2

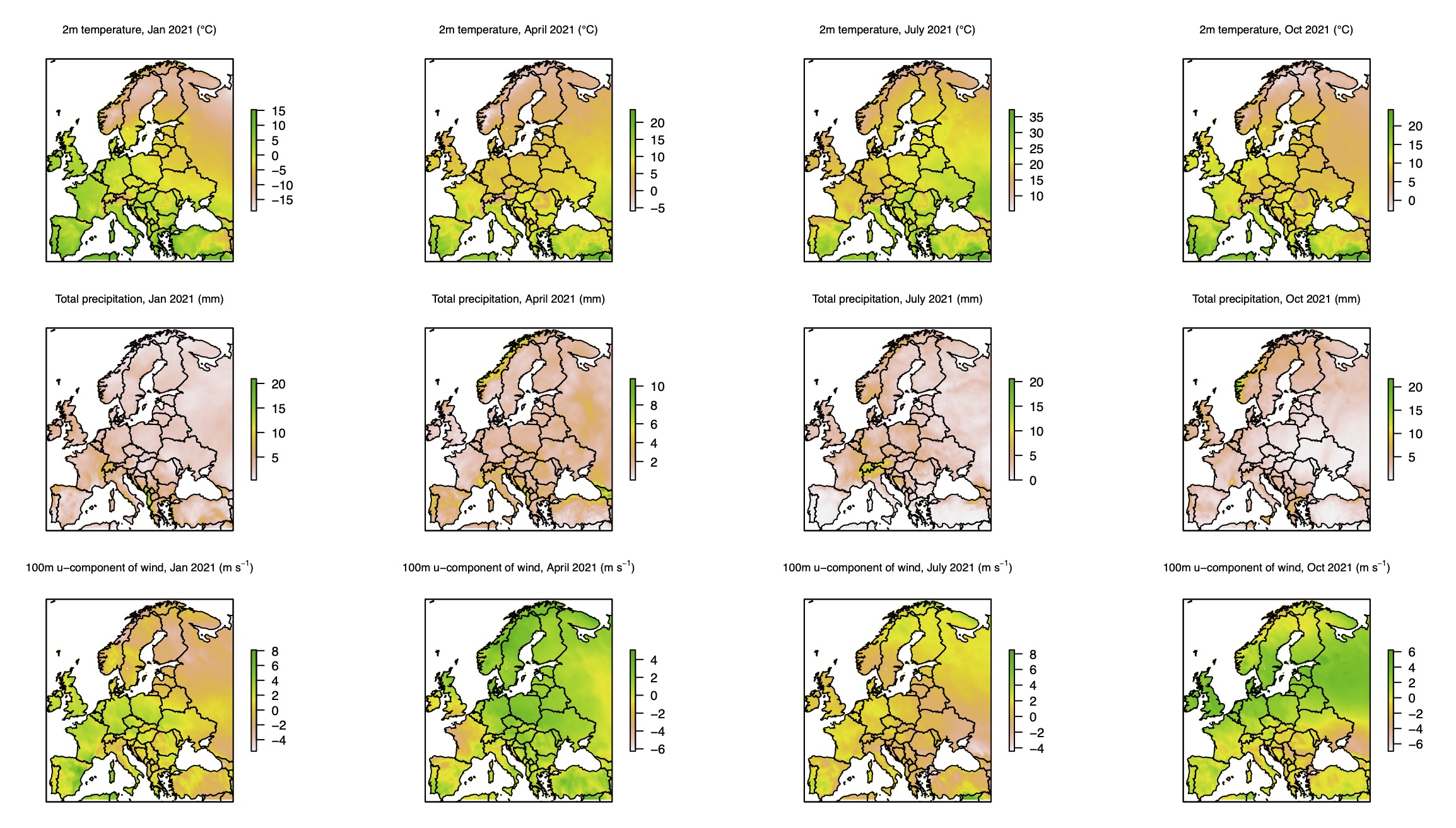

Figure S8 Distribution of predictors for viral genetic diversity of West Nile virus over time in Europe. Data in four months (January, April, July, and October) were shown for each predictor. Definition for each predictor is shown in Supporting file 3.

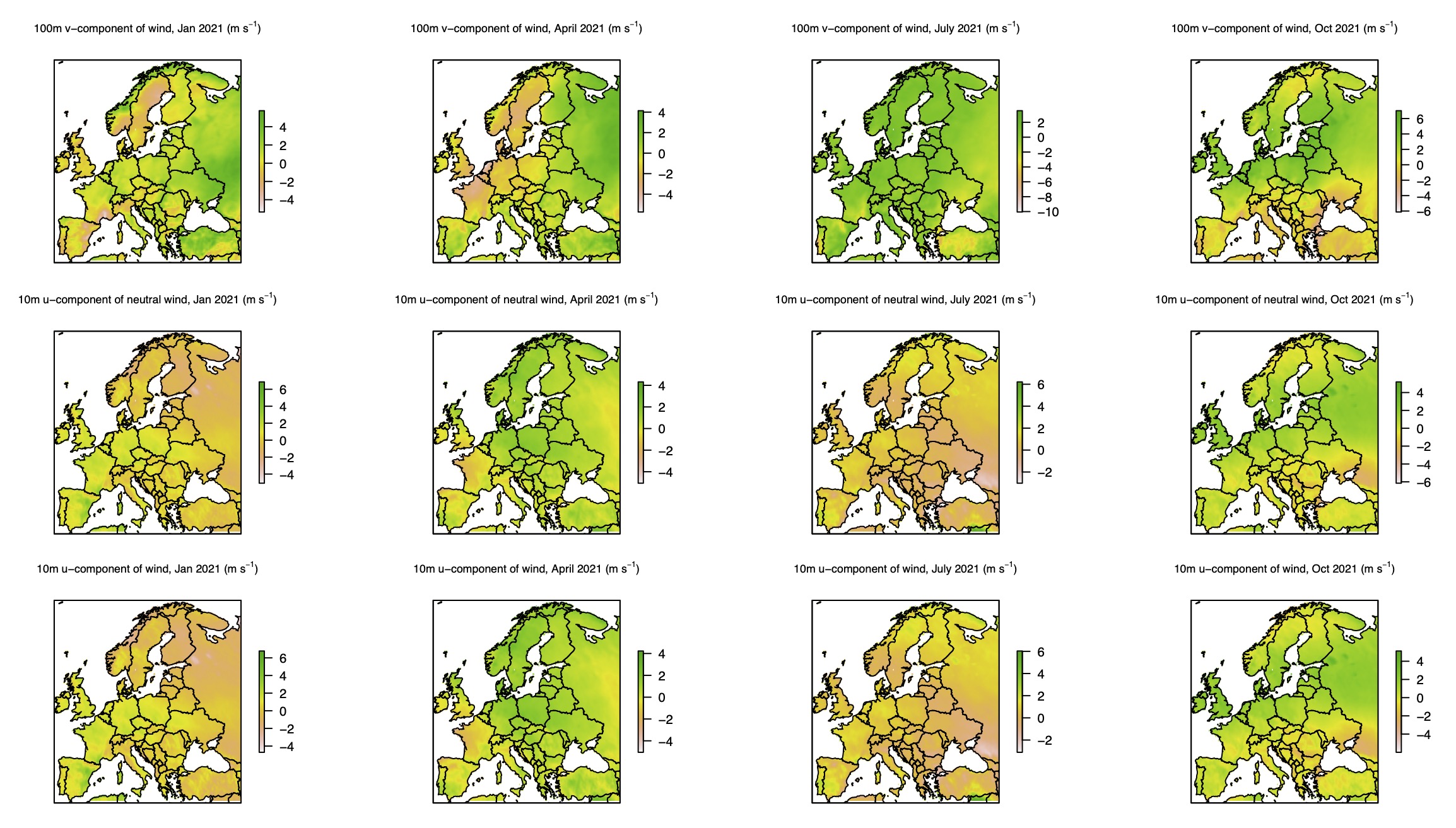

Figure S8 continued - 1

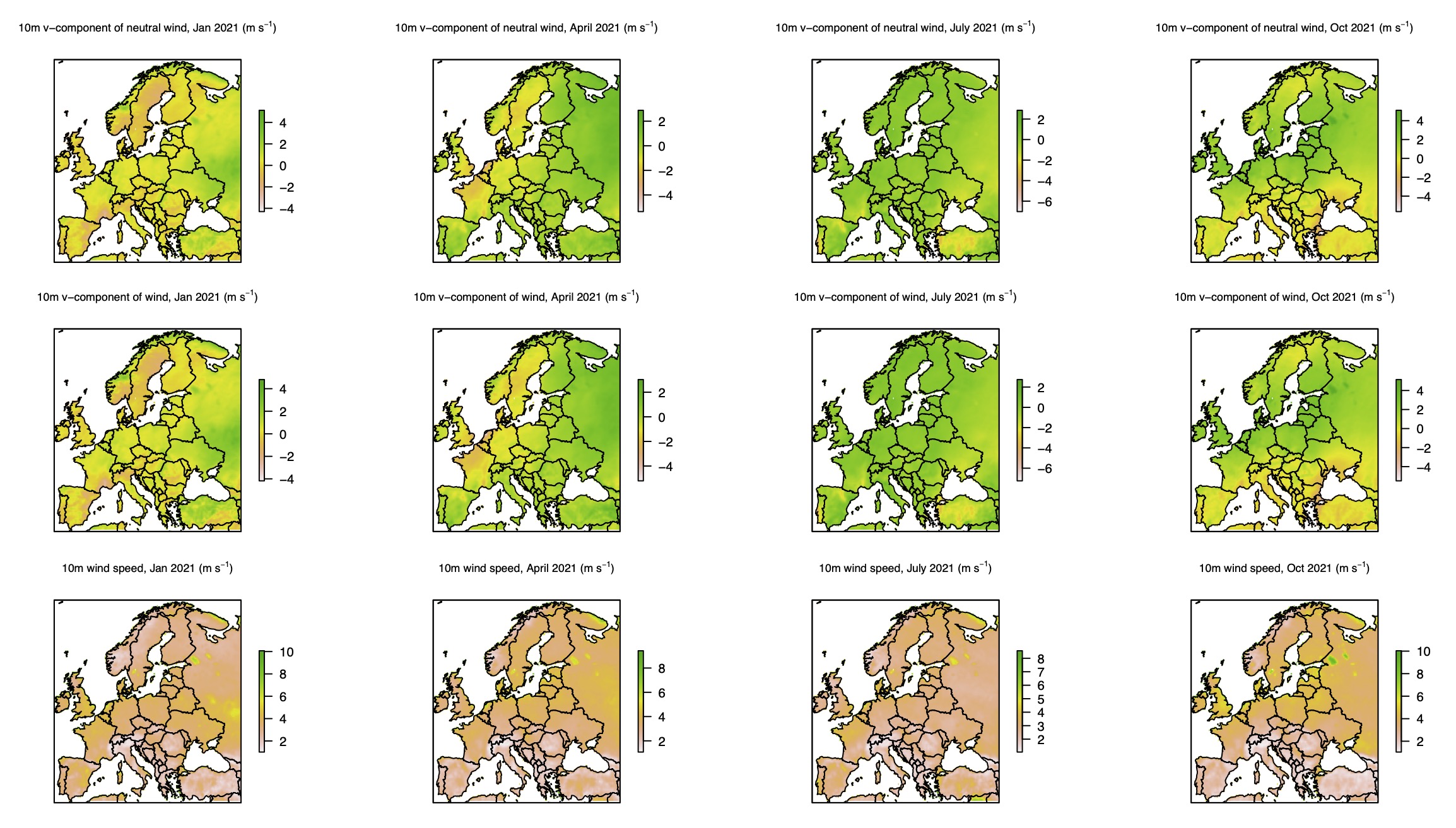

Figure S8 continued - 2

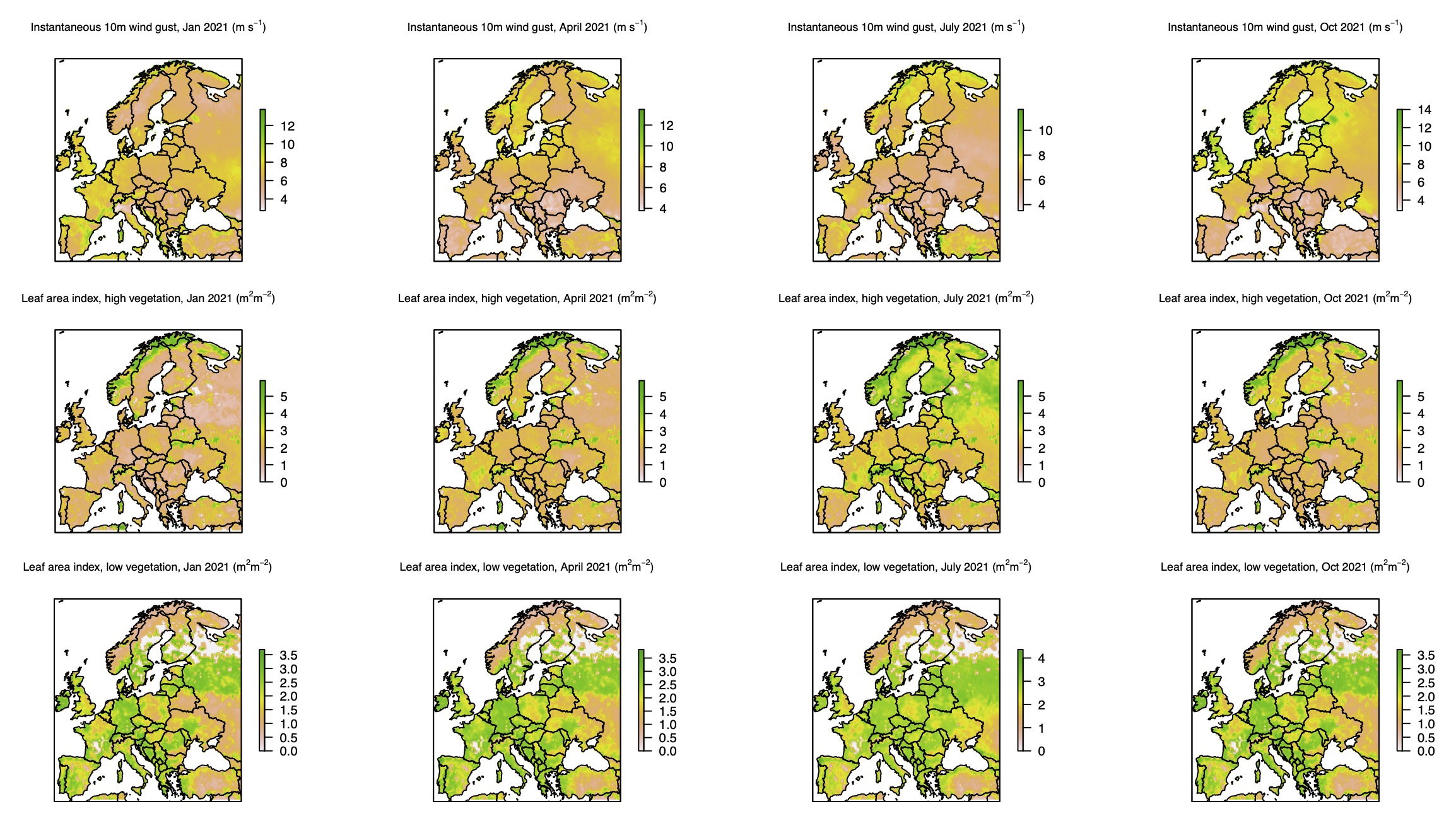

Figure S8 continued - 3

### Supplementary Movie

Movie S1 The movie of dispersal history of WNV-2a in Europe

### Supporting files

Supporting file 1 Sequences and metadata of WNV-2a used in this study

Supporting file 2 Predictors for dispersal of West Nile virus in Europe

Supporting file 3 Predictors for viral genetic diversity of West Nile virus over time in Europe
