## Supporting file 2 for "West Nile Virus spread in Europe - phylogeographic pattern analysis and key drivers"

Supporting file 2 Predictors for dispersal of West Nile virus in Europe assessed for quality according to the methodology of Horigan et al., 2022

| **Group** | **Predictor of interest** | **Definition** | **Original Resolution** | **Source** | **Accuracy & Precision** | **Reliability & Consistency** | **Timeliness** | **Completeness** | **Availability & Accessibility** | **Granularity** | **Total** |
| --- | --- | --- | --- | --- | --- | --- | --- | --- | --- | --- | --- |
| Climatic | Annual mean temperature | Annual mean temperature (℃) of 2021 is calculated from monthly average temperature of air at 2m above the surface of land, sea, or inland waters. | 0.25° | ERA5 ^1^ | 3 | 3 | 3 | 3 | 2 | 2 | 16 |
|  | Annual total precipitation | Annual total precipitation (mm) of 2021 is calculated by summing the monthly accumulated liquid and frozen water, comprising rain and snow, that falls to the Earth's surface. |  |  |  |  |  |  |  |  |  |
| Land Cover | Deciduous Broadleaf Trees | The percentage (%) of each of the land cover type (Deciduous Broadleaf Trees, Evergreen Broadleaf Trees, Evergreen/Deciduous Needleleaf Trees, Mixed/Other Trees, and Shrubs) in each grid cell (data released in 2014).  Mixed/Other Trees include mixed forest, woody savanna, and savanna. | 30" | EarthEnv ^2^ | 3 | 2 | 2 | 3 | 2 | 3 | 15 |
|  | Evergreen Broadleaf Trees |  |  |  |  |  |  |  |  |  |  |
|  | Evergreen/Deciduous Needleleaf Trees |  |  |  |  |  |  |  |  |  |  |
|  | Mixed/Other Trees |  |  |  |  |  |  |  |  |  |  |
|  | Shrubs |  |  |  |  |  |  |  |  |  |  |
|  | Cropland | The percentage (%) of each of the land cover type (Cropland, Pasture, Urban land, Primary land, and Secondary land) of 2015 in each grid cell.  Primary land is the natural vegetation (either forest or non-forest) that has never been impacted by human activities since 1700; Secondary land is the natural vegetation (either forest or non-forest) that is recovering from previous human disturbance. | 0.25° | Land-Use Harmonization (LUH2) ^3^ | 2 | 2 | 2 | 3 | 3 | 2 | 14 |
|  | Pasture |  |  |  |  |  |  |  |  |  |  |
|  | Urban land |  |  |  |  |  |  |  |  |  |  |
|  | Primary land |  |  |  |  |  |  |  |  |  |  |
|  | Secondary land |  |  |  |  |  |  |  |  |  |  |
|  | Urbanization of cropland | The percentage (%) of land area change from cropland/pasture/primary land/secondary land to urban land (Urbanization of cropland, Urbanization of pasture, Urbanization of primary land, and Urbanization of secondary land) in 2015 in each grid cell. |  |  |  |  |  |  |  |  |  |
|  | Urbanization of pasture |  |  |  |  |  |  |  |  |  |  |
|  | Urbanization of primary land |  |  |  |  |  |  |  |  |  |  |
|  | Urbanization of secondary land |  |  |  |  |  |  |  |  |  |  |
|  | Cultivated and Managed Vegetation | The percentage (%) of each of the vegetation type (Cultivated and Managed Vegetation, Herbaceous Vegetation, and Regularly Flooded Vegetation) in each grid cell (data released in 2014).  Cultivated and Managed Vegetation include cropland and the mixture of cropland and natural vegetation;  Herbaceous Vegetation represents the grasslands;  Regularly Flooded Vegetation represents the permanent wetlands. | 30" | EarthEnv ^2^ | 3 | 2 | 2 | 3 | 2 | 3 | 15 |
|  | Herbaceous Vegetation |  |  |  |  |  |  |  |  |  |  |
|  | Regularly Flooded Vegetation |  |  |  |  |  |  |  |  |  |  |
|  | Open water | The percentage (%) of open water in each grid cell (data released in 2014). |  |  |  |  |  |  |  |  |  |
|  | wetland_combine | Variable wetland comprises 12 different wetland types including lake, reservoir, river, freshwater/marsh/floodplain, swamp forest/flooded forest, coastal wetland (incl. mangrove, estuary, delta, lagoon), pan/brackish/saline wetland, bog/fen/mire (peatland), intermittent wetland/lake, 50-100% wetland, 25-50% wetland, and wetland complex (0-25% wetland) (data released in 2004). We rescaled wetland as a binary variable: presence or absence of wetland.  For wetland_combine, we included all 12 ranges comprising lakes, reservoirs, rivers and different wetland types. | 30" | Global Lakes and Wetlands Database (GLWD) ^4^ | 2 | 2 | 2 | 3 | 2 | 3 | 14 |
|  | lake_river_reservoir | Range 1 to 3 comprises lakes, reservoirs, rivers |  |  |  |  |  |  |  |  |  |
|  | wetland_other | Range 4 to 12 comprises other different wetland types. |  |  |  |  |  |  |  |  |  |
|  | Wetland concentration | Coastal wetland intensity in Europe in 2000, with special interest in the coastal wetlands. It ranges 0 to 100. | 30" | Wetland concentration in Europe (2000) ^5^ | 2 | 2 | 2 | 2 | 3 | 3 | 14 |
| Topography | Elevation | Land surface elevation (m) of 2010 in each grid cell | 0.1° | Global Multi-resolution Terrain Elevation Data (GMTED) 2010 dataset ^6^ | 2 | 3 | 2 | 3 | 3 | 3 | 16 |
| Socio-economic | Human population | Human population count of 2020 (counts, in persons) | 0.25° | Gridded Population of the World, Version 4 (GPWv4) ^7^ | 2 | 2 | 2 | 3 | 2 | 3 | 14 |
|  | Gross domestic product (GDP) | GDP (PPP, 2011 int. USD) of 2015 | 5' | Gridded global datasets for Gross Domestic Product and Human Development Index ^8^ | 2 | 2 | 2 | 3 | 3 | 3 | 15 |
| Biodiversity | Livestock count | Domestic animal headcount, summed sheep, pig, horse, goat, cattle, and buffalo of 2010 (counts) | 5' | Gridded Livestock of the World (GLW 3) ^9^ | 2 | 2 | 1 | 2 | 3 | 3 | 13 |
|  | Mammal species richness | Aggregation of the presence grids data for the entire class (counts, data released in 2015) | 30" | Global Mammal Richness Grids ^10^ | 2 | 2 | 2 | 3 | 2 | 3 | 14 |
|  | Flyway_Anseriformes | Flyway of 3 bird groups (mainly in European countries), including Anseriformes (White-fronted goose and Barnacle geese), Apodiformes (Apus melba), and Passeriformes (Red-backed shrike and Whinchats Saxicola rubetra) | NA | Movebank for animal tracking data ^11^ | 2 | 2 | 2 | 2 | 3 | 3 | 14 |
|  | Flyway_ Apodiformes |  |  |  |  |  |  |  |  |  |  |
|  | Flyway_Passeriformes |  |  |  |  |  |  |  |  |  |  |
|  | Birds Directive | This indicator, Birds and Habitats Directives, includes sites designated under the Birds Directive (Special Protection Areas/SPAs) and the Habitats Directive (Sites of Community Importance/SCIs, and Special Areas of Conservation/SACs) in 2000. We treat this indicator as three binary predictors, i.e., presence and absence of birds directives, habitats directives, as well as both birds and habitats directives. | Polygon | Natura 2000 data ^12^ | 2 | 2 | 2 | 3 | 3 | 3 | 15 |
|  | Habitats Directive |  |  |  |  |  |  |  |  |  |  |
|  | Birds and Habitats directive |  |  |  |  |  |  |  |  |  |  |
|  | Richness of forest-related species and habitats | This dataset refers to the Richness index of Species and Habitats of Conservation Concern indicator in 2012. It is composed itself of three sub-indicators: “Forest Non-bird species”, “Forest bird species” and “Forest habitats”. The sub-indicators were then normalized for each European forest type and successively combined not assigning any specific weight to a particular sub-indicator. This indicator ranges between 0 and 1. | 30" | Richness of forest-related species and habitats indicator 2012 dataset ^13^ | 1 | 2 | 2 | 2 | 3 | 3 | 13 |
|  | Culex pipiens status | The current known distribution of the Culex pipiens group (Culex pipiens and Culex torrentium) in Europe in September 2021. | Regional level | Culex pipiens group - current known distribution ^14^ | 2 | 3 | 2 | 1 | 3 | 1 | 12 |

NA, not applicable

**Reference**

1. Hersbach H, Bell B, Berrisford P, et al. ERA5 monthly averaged data on single levels from 1959 to present: Copernicus Climate Change Service (C3S) Climate Data Store (CDS), 2019.

2. Tuanmu M-N, Jetz W. A global 1-km consensus land-cover product for biodiversity and ecosystem modelling. *Global Ecology and Biogeography* 2014;23(9):1031-45. doi: doi:10.1111/geb.12182

3. Hurtt GC, Chini L, Sahajpal R, et al. Harmonization of global land use change and management for the period 850–2100 (LUH2) for CMIP6. *Geosci Model Dev* 2020;13(11):5425-64. doi: 10.5194/gmd-13-5425-2020

4. World Wildlife Fund - WWF. Global Lakes and Wetlands Database: Lakes and Wetlands Grid (Level 3), 2004.

5. European Environment Agency - EEA. Wetland concentration in Europe (2000), 2009.

6. Amatulli G, Domisch S, Tuanmu M-N, et al. A suite of global, cross-scale topographic variables for environmental and biodiversity modeling. *Scientific Data* 2018;5(1):180040. doi: 10.1038/sdata.2018.40

7. Center for International Earth Science Information Network-CIESIN-Columbia University. Gridded Population of the World, Version 4 (GPWv4): Population Count, Revision 11. Palisades, New York: NASA Socioeconomic Data and Applications Center (SEDAC), 2018.

8. Kummu M, Taka M, Guillaume J. Gridded global datasets for Gross Domestic Product and Human Development Index over 1990-2015: Dryad, 2020.

9. Gilbert M, Nicolas G, Cinardi G, et al. Global distribution data for cattle, buffaloes, horses, sheep, goats, pigs, chickens and ducks in 2010. *Scientific Data* 2018;5(1):180227. doi: 10.1038/sdata.2018.227

10. International Union for Conservation of Nature - IUCN., Center for International Earth Science Information Network - CIESIN - Columbia University. Gridded Species Distribution: Global Mammal Richness Grids, 2015 Release. Palisades, New York: NASA Socioeconomic Data and Applications Center (SEDAC), 2015.

11. Max Planck Institute of Animal Behavior, North Carolina Museum of Natural Sciences, University of Konstanz. Movebank for animal tracking data, 2021.

12. European Environment Agency - EEA. Natura 2000 data - the European network of protected sites, 2020.

13. European Environment Agency - EEA. Richness of forest-related species and habitats indicator 2012 dataset, 2012.

14. European Centre for Disease Prevention and Control - ECDC. Culex pipiens group - current known distribution: September 2021, 2021.
