## Supporting file 3 for "West Nile Virus spread in Europe - phylogeographic pattern analysis and key drivers"

Supporting file 3 Predictors for viral genetic diversity of West Nile virus over time in Europe, 2004-2021

| **Predictor of interest** * | **Definition from the data source** |
| --- | --- |
| Surface air temperature at 2m | Monthly temperature (℃): the temperature of air at 2m above the surface of land, sea or inland waters. |
| Precipitation | Monthly precipitation (mm): the accumulated liquid and frozen water, comprising rain and snow, that falls to the Earth's surface. |
| 100m u-component of wind | This parameter is the eastward component of the 100 m wind (m s^-1^). It is the horizontal speed of air moving towards the east, at a height of 100 metres above the surface of the Earth. |
| 100m v-component of wind | This parameter is the northward component of the 100 m wind (m s^-1^). It is the horizontal speed of air moving towards the north, at a height of 100 metres above the surface of the Earth. |
| 10m u-component of neutral wind | This parameter is the eastward component of the "neutral wind"(m s^-1^), at a height of 10 metres above the surface of the Earth. The neutral wind is calculated from the surface stress and the corresponding roughness length by assuming that the air is neutrally stratified. The neutral wind is slower than the actual wind in stable conditions, and faster in unstable conditions. The neutral wind is, by definition, in the direction of the surface stress. The size of the roughness length depends on land surface properties or the sea state. |
| 10m v-component of neutral wind | This parameter is the northward component of the "neutral wind" (m s^-1^), at a height of 10 metres above the surface of the Earth. Detailed information on the “neutral wind” has been described above. |
| 10m u-component of wind | This parameter is the eastward component of the 10m wind (m s^-1^). It is the horizontal speed of air moving towards the east, at a height of ten metres above the surface of the Earth. |
| 10m v-component of wind | This parameter is the northward component of the 10m wind (m s^-1^). It is the horizontal speed of air moving towards the north, at a height of ten metres above the surface of the Earth. |
| 10m wind speed | This parameter is the horizontal speed of the wind, or movement of air (m s^-1^), at a height of ten metres above the surface of the Earth. The eastward and northward components of the horizontal wind at 10m are also available as parameters. |
| Instantaneous 10m wind | This parameter is the maximum wind gust (m s^-1^) at the specified time, at a height of ten metres above the surface of the Earth. The WMO defines a wind gust as the maximum of the wind averaged over 3 second intervals. This duration is shorter than a model time step, and so the ECMWF Integrated Forecasting System (IFS) deduces the magnitude of a gust within each time step from the time-step-averaged surface stress, surface friction, wind shear and stability. |
| Leaf area index, high vegetation | This parameter is the surface area of one side of all the leaves found over an area of land for vegetation classified as "high" (m^2^ m^-2^). "High vegetation" consists of evergreen trees, deciduous trees, mixed forest/woodland, and interrupted forest. This parameter has a value of 0 over bare ground or where there are no leaves. |
| Leaf area index, low vegetation | This parameter is the surface area of one side of all the leaves found over an area of land for vegetation classified as "low"(m^2^ m^-2^). "Low vegetation" consists of crops and mixed farming, irrigated crops, short grass, tall grass, tundra, semidesert, bogs and marshes, evergreen shrubs, deciduous shrubs, and water and land mixtures. Similarly, this parameter has a value of 0 over bare ground or where there are no leaves |
| European common bird index | The population changes index of the 170 common bird species in Europe between 1980 to 2019 provided by PanEuropean Common Bird Monitoring Scheme (PECBMS) (https://pecbms.info/trends-and-indicators/). Data is classified into three groups (farmland, forest and others) and classified into different bird orders (*Galliformes, Passeriformes, Strigiformes, Accipitriformes, Galliformes, Coliformes, and Falconiformes*) and tested in our study. |

* all data except bird index and human cases in this table had an original resolution of 0.25° and were all obtained from ERA5 ^1^.

Reference

1. Hersbach H, Bell B, Berrisford P, et al. ERA5 monthly averaged data on single levels from 1959 to present: Copernicus Climate Change Service (C3S) Climate Data Store (CDS), 2019.
