## Supplementary figures and images for "West Nile Virus spread in Europe - phylogeographic pattern analysis and key drivers"

### Movie S1

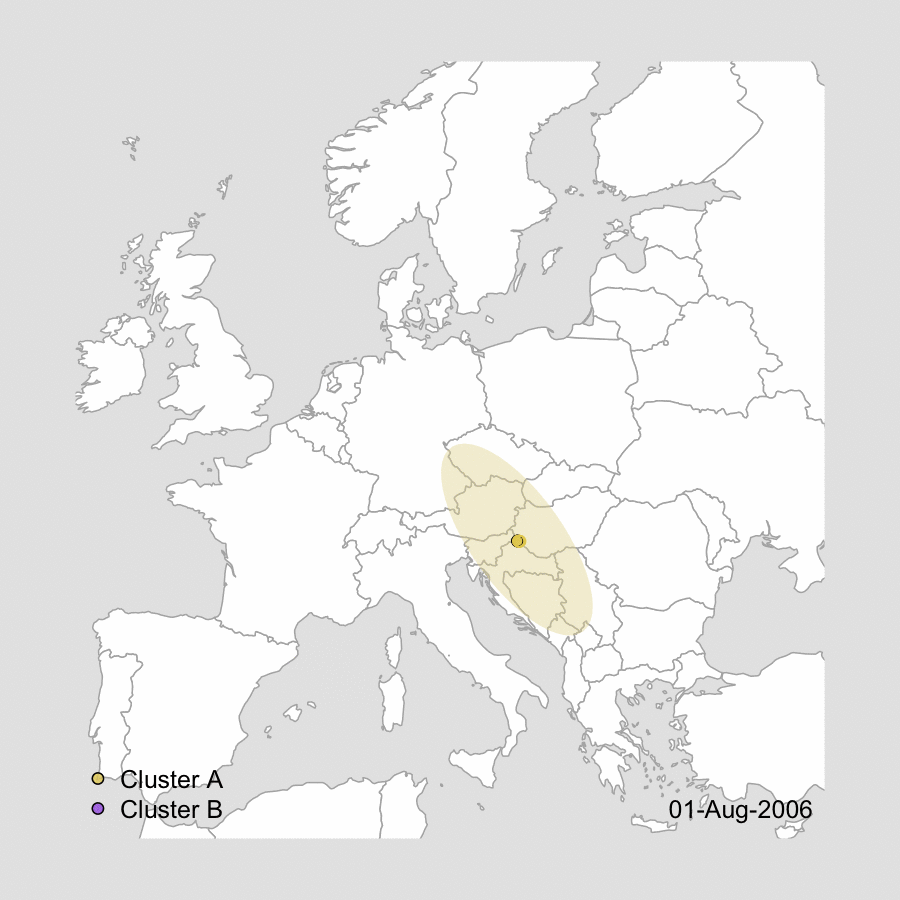
